# MIL Cell – A tool for multi-scale simulation of yeast replication and prion transmission

**DOI:** 10.1101/2023.03.21.533288

**Authors:** Damien Hall

## Abstract

The single celled baker’s yeast, *Saccharomyces cerevisiae*, can sustain a number of amyloid-based prions, with the three most prominent examples being [URE3] formed from the Ure2 protein (a regulator of nitrogen catabolism), [PSI+] formed from the Sup35 protein (a yeast translation termination release factor) and [PIN+] formed from the Rnq1 protein (of as yet unknown function). In a laboratory environment, haploid *S. cerevisiae* cells of a single mating type can acquire an amyloid prion in one of two ways (i.) Spontaneous nucleation of the prion within the yeast cell, and (ii.) Receipt via mother-to-daughter transmission during the cell division cycle. Similarly, prions can be lost from a yeast due to (i) Dissolution of the prion amyloid by its breakage into non-amyloid monomeric units, or (ii) Preferential donation/retention of prions between the mother and daughter during cell division. Here we present a computational tool, called MIL-CELL, for modelling these four general processes using a multiscale approach that is able to describe both spatial and kinetic aspects of the yeast life cycle and the amyloid- prion behavior. The yeast growth cycle is considered in two stages, a mature yeast that is competent to bud (M), and a daughter yeast (D) defined as a fully grown and detached bud. In the virtual plate experiment each transition in yeast growth is stochastically regulated, according to temporal and spatial characteristics, in a manner able to incorporate concepts of confluent growth. Between the relatively coarse time-points used for the particle level description, a set of differential equations, describing the nucleation, growth, fragmentation and clumping of amyloid fibrils, is solved numerically, for each individual yeast cell. Distribution of amyloid between the mother and the daughter is carried out by solving a set of kinetic partition equations between mother and the newly forming (and still attached) daughter during the yeast budding stage. In this paper we describe the workings of the model, the assumptions upon which it is based and some interesting simulation results that pertain to wave-like spread of the epigenetic prion elements through the yeast population. MIL-CELL (**M**onitoring **I**nduction and **L**oss of prions in **Cell**s) is provided as a stand-alone graphical user interface-based executable program for free download with the paper (supplementary section).

**MIL-CELL download:**

https://drive.google.com/drive/folders/1xNBSL_2sGNkyXfYLYUyXjyM9ibGAcQUL?usp=sharing

## Introduction

The first yeast epigenetic factor to be identified as an amyloid prion was [URE3], which is assembled via polymerization of the Ure2p protein [**Wickner, 1994; Masison et al 1997; King et al. 1997; Wickner et al. 1999**]. Since that time numerous other yeast amyloid prions have been discovered, with the two most notable examples being [PSI+] (generated from the Sup35 protein) [**Wickner, 1994; Patino et al. 1996; Paushkin et al. 1996; Derkatsch et al. 1996**] and [PIN+] (assembled from the Rnq1 protein) **[Derkatsch et al. 1996; Derkatsch et al. 2000; Sondheimer and Lindquist, 2000]**. Relatively recently, researchers have taken up the challenge of producing biophysical models of amyloid growth in yeast **[Tanaka et al. 2006; Lemarre et al. 2020; Banwarth-Kuhn and Sindi, 2020**]. Whilst successful in their specified aims, these models have neglected certain important physical aspects related to the (i) effects of spatial arrangement of the growing cells within the colony on the dispersion of amyloid amongst the yeast, (ii) biochemical mechanism of amyloid growth and transfer between yeast and nascent daughter, and (iii) the biochemical and physical determinants of the colony screen. The aim of the current work was to develop an informative biophysical model of amyloid formation and cytosolic transfer in dividing yeast that could usefully comment on these previously neglected features [**Tanaka et al. 2006; Lemarre et al. 2020; Banwarth-Kuhn and Sindi, 2020**]. To help orient the reader, in what follows, we first provide a short history of the study of amyloid prion growth and transmission in yeast. After setting this introductory foundation we develop the theoretical basis of our model and then use it to simulate some interesting situations of yeast carrying and passing on amyloid prions to their offspring. We conclude with a short description of the usage of the MIL-CELL program and describe some of its potential future applications.

### A short history of amyloid/prions in Saccharomyces cerevisiae

Although the single celled baker’s yeast *Saccharomyces cerevisiae* is one of the simplest eukaryotic organisms, it shares many genetic and biochemical pathways in common with more complex eukaryotic organisms (including humans), and for this reason it has become a key model system [**Karathia et al. 2011; Duina et al. 2015**]. Despite being regarded as relatively simple, *S. cerevisiae* nevertheless, exhibits a complex life-cycle, that is capable of mitotic reproduction from two vegetative states of different ploidy (yeast budding from both the haploid and diploid^1^ states), meiotic cell division (yeast sporulation) from its diploid form, and sexual reproduction (yeast mating) between the different sexes (*a* and *α*) of haploid yeast states [**Duina et al. 2015**] (**Fig. 1**). Due to its approximately 90-minute reproduction time and an abundance of yeast specific biochemical and genetic experimental tools, *S. cerevisiae*, has been pivotal to the development of our modern scientific understanding of the eukaryotic cell cycle^2^ [**Hartwell, 1974; Hartwell and Unger, 1977; Forsburg and Nurse, 1991**]. From a number of somewhat initially confounding genetic studies a series of epigenetic factors^3^ were identified in yeast [**Riggs et al. 1996; Bonasio et al. 2010**] and the study of the non-chromosomal DNA sequence-related origins of such epigenetic factors helped to spawn important fields of research such as DNA methylation^4^ [**Singal and Ginder, 1999; Weissbach, 2013**], histone post-translational modification [**Davie et al. 1981; O’Kane and Hyland, 2019**], transposon biology [**Zou et al. 1996; Hosaka and Kakutani, 2018**], mitochondrial gene replication [**Rasmussen, 2003**] and the cytosolic localization of both dsRNA virus-like genomes [**Wickner and Leibowitz, 1977**] and yeast DNA plasmids [**Gunge et al. 1983**]. Within such a diverse background of non-chromosomal DNA sequence-based epigenetic factors there were two particular phenotypic traits, [URE3] – associated with catabolism of uredosuccinic acid as a potential yeast food source [**Lacroute 1971; Aigle and Lacroute, 1975**], and [PSI+] – associated with the suppression of nonsense genes produced by translation past a stop codon [**Cox 1965; Serio et al. 1999**], which proved enigmatic and resisted easy assignment to any of the above noted epigenetic causes [**Tuite et al. 2015**].

**Figure 1:**
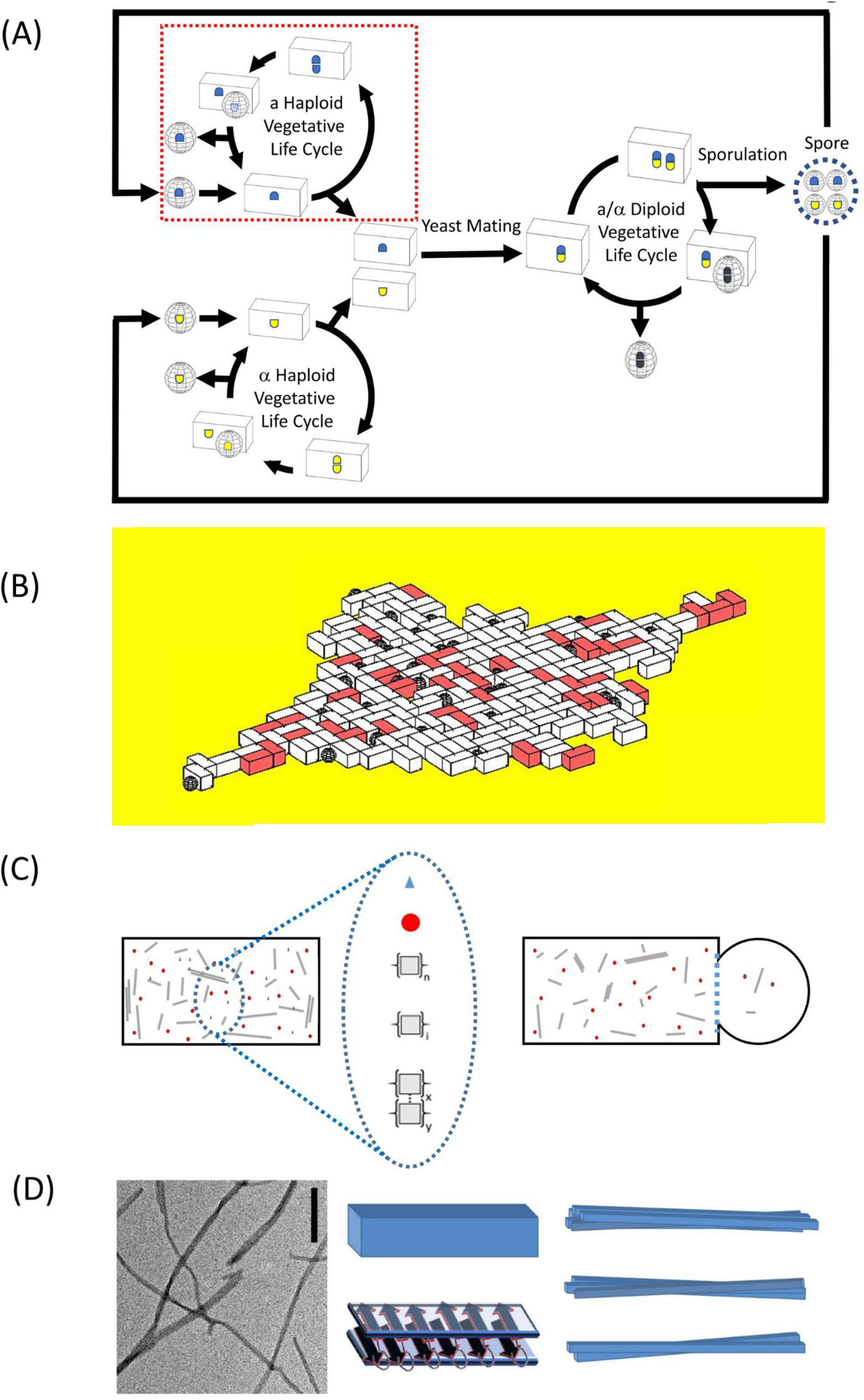
An overview of areas covered by the MIL- CELL program. **(A) Complex yeast life cycle:** *Saccharomyces Cerevisiae* is capable of mitotic reproduction from two vegetative states of different ploidy (yeast budding from both the haploid and diploid states), meiotic cell division (yeast sporulation) from its diploid form, and sexual reproduction (yeast mating) between different sex (a and α) haploid yeast states. This work considers only the yeast haploid vegetative life cycle (highlighted in red dotted box). **(B) Particle modelling of the growth and division of haploid yeast:** The haploid yeast growth/division is simulated in two dimensions using a stochastic particle model sensitive to both intrinsic growth rates and the density of surrounding yeast. The presence or absence of yeast prions is indicated using a color scale that corresponds to the biochemical color development assay applied experimentally. **(C) Chemical modelling of the growth and transfer of cytosolic amyloid ‘prions’ within yeast:** The dynamical growth and partition behavior of the cytosolic prion components is modelled using a set of partial differential equations. (Left) Schematic showing a snapshot of the yeast cell’s cytosolic contents and their relative concentrations (blue triangle - amino acids; red circle - amyloid monomer; n linked squares - amyloid nucleus; i linked squares- amyloid protofibril; laterally aligned squares- clumped fibers). (Right) Schematic showing the transfer of amyloid prions between mother and daughter cells during yeast division. **(D) Insight into amyloid structure:** The [PSI+} phenotype is conferred by the presence of amyloid prions formed from homo-polymerization of the Sup35 protein. (Left) Typical transmission electron micrograph (TEM) image of amyloid fibers (scale bar 100 nm). (Middle) Amyloid fibers are typically long and thin with a length distribution in the range of nm to μm and width distribution in the range 4–20 nm. Single amyloid proto-filaments are formed as a result of polypeptide units forming intermolecular β-sheets along the long axis of the fiber. The example diagram shows a rectangular box representation of a protofilament which is formed by a polypeptide with two stacks of β-sheet. (Right) Amyloid proto-filaments can undergo self-association to form clumped fibers also called ‘mature fibrils’ which are typically helical or lateral arrangements of multiple protofilaments. See or lateral arrangements of multiple protofilaments. See **[Hall and Edskes, 2012]**.

Adapting concepts developed in the field of Scrapie biology [**Prusiner, 1982**], Reed Wickner proposed a paradigm shifting ‘protein only’ epigenetic mechanism for the [URE3] phenotype that involved prion amyloid formation from the Ure2p protein ^5^ [**Wickner 1994**]. Wickner’s proposal was based on a set of experiments that involved overproduction of Ure2p, cytoplasmic transfer via cytoduction and reversible cycles of losing/regaining the [URE3] phenotype^6^ [**Wickner 1994; Wickner et al. 1995; Masison et al. 1997; King et al. 1997; Edskes et al. 1999; Wickner et al. 1999**]. In his original paper [**Wickner, 1994**], Wickner additionally suggested that an amyloid-based mechanism, involving aggregation of the Sup35 protein^7^, would also be consistent with experimental knowledge concerning the [PSI+] yeast phenotype [**Cox 1965; Tuite et al. 1983; Doel et al. 1994**]. The [PSI+] growth phenotype, which was originally discovered by Cox in 1965 [**Cox, 1965**], is now known to result from an unusually high production of translational read through events^8^ [**Didichenko et al. 1991; Stansfield et al. 1995; Serio et al. 1999**]. Following Wickner’s suggestion of its potential amyloid prion nature, a range of genetic [**Patino et al. 1996**] and biochemical investigations into [PSI+] (that even involved introduction of an external amyloid created in vitro from recombinantly synthesized Sup35 protein into yeast to induce a stable [PSI+] phenotype) confirmed the amyloid prion basis of [PSI+] inheritance [**Sparrer et al 2000**]. A third major amyloid prion system (in terms of applied research effort), termed [PIN+], was later discovered **[Derkatsch et al. 1997; Sondheimer and Lindquist, 2000**]. Derived from the acronym for [PSI+] Inducibility, the [PIN+] phenotype was identified as an additional requirement for the production of [PSI+] amyloid (and the associated [PSI+] associated phenotypic traits) [**Derkatch et al 2000; Sondheimer and Lindquist, 2000**]. Using similar genetic and biochemical procedures the [PIN+] epigenetic trait was shown to be due to conversion of the Rnq1 protein into amyloid form [**Patel and Liebman, 2007**].

Since Wickner’s original discovery that certain non-chromosomal epigenetic traits in yeast could be effected by an amyloid prion mechanism [**Wickner 1994**] a range of additional yeast prions have been discovered **[Wickner et al. 2015]**. However, the question as to whether yeast prions represent a disease or a potential benefit to yeast is still debated [**Wickner et al. 2011; Halfmann et al. 2012; Wang et al. 2017**]. In a manner that both precedes and runs parallel to, the discovery of the amyloid basis of the yeast epigenetic factors [URE3], [PSI+] and [PIN+], our general understanding of amyloid structure [**Glenner et al. 1974; Lansbury et al. 1992; Adamcik et al. 2010; Jahn et al. 2012; Hall 2012; Eisenberg and Sawaya, 2017; Meier et al. 2017; Iadanza et al. 2018]**, its mechanism of formation [**Masel et al. 1999; Pallito and Murphy, 2001**; **Hall and Edskes, 2004; Hall et al., 2015; Hirota et al. 2019**] and its negative associations with the set of devastating amyloidosis diseases [**Glenner and Wong, 1984; Nowak et al. 1999; Merlini and Bellotti, 2003**; **Hall and Edskes, 2009, 2012; Martinez-Naharro et al. 2018; Weickenmeier et al. 2018; Fornari et al. 2019**] has continued apace. Since its original identification from patient biopsy/autopsy at the macroscopic [for an early history see **Sipe and Cohen, 2000**] and molecular levels [**Cohen and Calkins, 1959; Bladen et al. 1966; Eanes and Glenner, 1968; Prusiner et al. 1983; Glenner and Wong, 1984**] our present-day collective understanding of the disastrous consequences arising from defects in the biological control systems regulating protein folding and amyloid production in vivo [**Hardy and Higgins, 1992; Labbadia and Morimoto, 2015; Klaips et al. 2018**] means that work directed at both delineating, and potentially controlling, the factors affecting these processes is, without hyperbola, of the utmost importance [**Ohtsuka and Suzuki, 2000; Aguzzi and Sigurdson, 2004; Ringe and Petsko, 2009; Wentink et al. 2019**]. Due to yeast possessing many of the same genes and proteins as those found in humans, the *S. cerevisiae* model system presents itself as an ideal vehicle for interrogation of biological factors affecting amyloid growth within a biological setting. Combining molecular biology and yeast genetic methods provides the experimenter with the ability to add (knock in) or remove (knockout) genes, switch particular genes on or off (silence or enhance expression) and, in some cases, to subtly tune the expression levels of particular proteins such as those associated with yeast chaperone and vacuole^9^ systems, to control the induction and loss of amyloids within an in vivo setting [**Chernova et al. 2017; Son and Wickner, 2019; Wickner et al. 2021**]. However, due to the highly complex and potentially non-linear nature of amyloid growth and transfer within a dividing and expanding set of cells, the results of such experiments crucially require simplifying (but not simple) mathematical models to aid with their interpretation. It is towards this goal that the present work is directed.

#### How does the MIL-CELL computational tool work?

To simulate the growth and transmission of amyloid prion elements within and between a population of yeast cells we have developed a multiscale modelling approach called MIL-CELL (with this acronym standing for Monitoring Induction and Loss of prions in Cells). In our approach the division and growth of yeast is described at the particle level whilst the behaviour of the amyloid prion elements is described microscopically using a set of chemical rate equations. In the next sections we describe these two approaches in turn before then explaining how they are coupled together. The simulation format is designed to match with a particular type of yeast culture experiment in which cells are grown and monitored at one layer thickness under a coverslip **[Cerulus et al. 2016; Mayhew et al. 2017; Zhao et al. 2018]** or via microfluidic/cell sorting assay [**Scheper et al 1987; Huberts et al. 2013**] thus reducing the problem to one of growth in either zero or two spatial dimensions.

### (i)#Particle level model of yeast life cycle

#### Factors determining the growth and division of yeast within a colony

As per **Fig. 2 (Top Panel)** we consider two distinct yeast states, a mature mother (M) and an immature daughter (D) [**Hartwell and Unger, 1977; Cerulus et al. 2016**]. These two states respectively undergo the following two transitions within the yeast life cycle (i.) Mother yeast producing a daughter yeast (**Eqn. 1a**), and (ii.) Daughter cell growing into a mother yeast (**Eqn. 1b**). Mature mother yeast are physically modelled as rectangular solids of aspect ratio 2 with actual dimensions of length, width and height respectively given by L_M_ = 5 μm, W_M_ = 2.5 μm and H_M_ = 2.5μm **[Nagel, 1946; Hartwell and Unger, 1977; Cerulus et al. 2016]**. Immature yeast are represented as spheres with a radius, R_D_, equal to half the width of the mature cell i.e. R_D_ = 1.25 μm. At time zero, a single mature yeast cell is placed on a central area of a plate defined by a sector wedge, beyond the bounds of which the yeast is unable to grow. Progression through the yeast life-cycle is modelled as a series of transitions, with each advance governed by a pairing of a minimum time delay, δ, and a transition time, τ specified in relation to a time, t, recorded from the starting point of its previous transition (**Eqn. 1**).

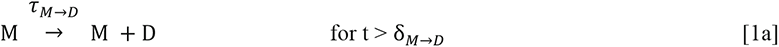

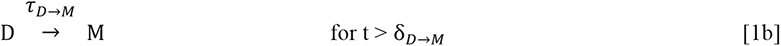

**Figure 2:**
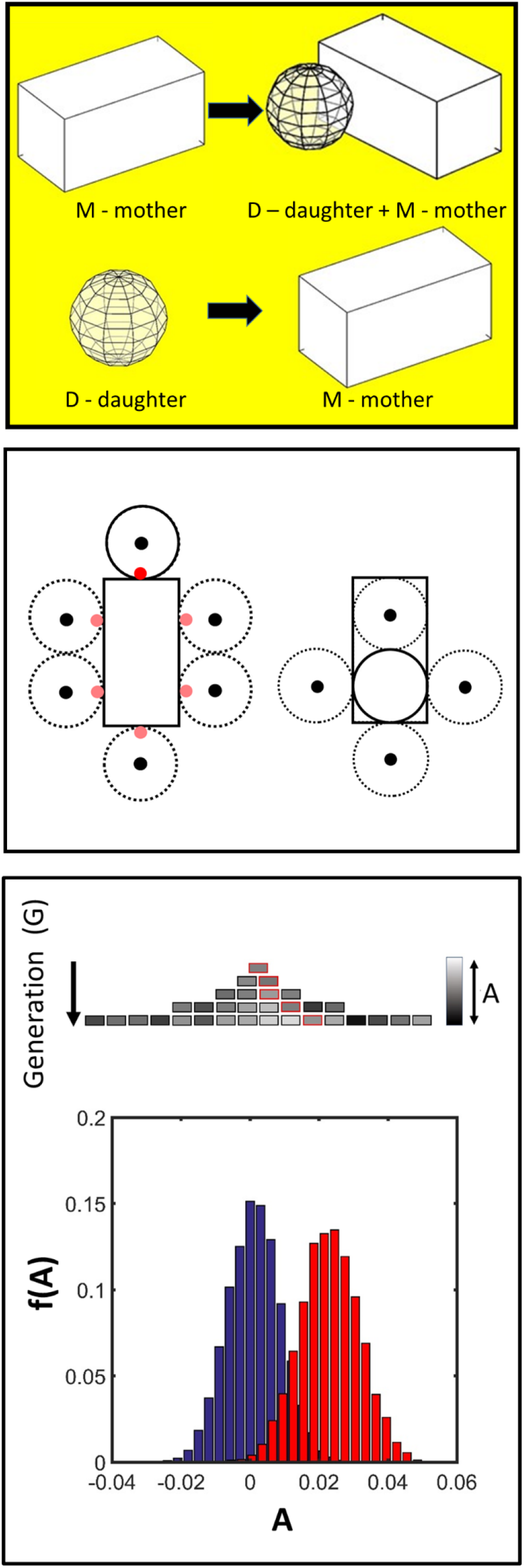
Considerations within the particle-level model of yeast growth. **(Top Panel) Schematic of yeast particle model:** Two different stable states of S. cerevisiae are considered, the yeast daughter (D) (modelled as a sphere) and the yeast mother (M) (modelled as a rectangular solid). **Two transitive states are considered (Top)** Mother yeast producing a daughter (M → M + D) (Eqn 1a). **(Bottom)** Daughter yeast growing to become a potential mother yeast (D → M) (Eqn. 1b). **(Central Panel) Schematic (shown in two dimensions) of the two types of S. cerevisiae cellular communal growth transitions considered: (Left) Mother to daughter transition** – An existing mother yeast (solid rectangle) is considered potentially able to form a bud at one of six equidistant locations in the horizontal plane (red circles). One of these six positions is randomly selected. If the stochastic sampling of the temporal (kinetic transition Eqn. 2b) and spatial (physical occupancy Eqn. 2c) criteria are met then growth of the daughter (solid circle) can occur (Eqn. 2a). If the stochastic sampling criteria are not met then a new position is selected without replacement and the process repeated. If all positions have been trialled without success then the growth process is considered unsuccessful **(Right) Daughter to potential mother transition** – An existing daughter (solid circle) may potentially expand to one of four equidistant positions filling a volume defining a rectangular solid (small black circles) if the stochastic sampling criteria of the temporal (Eqn. 2b) and spatial (Eqn. 2c) conditions are met. As per the mother to daughter transition a potential growth position is selected randomly and the testing performed without replacement until either a successful growth event is recorded to form a potential mother cell (solid rectangle) or all growth options have been rejected. For both transition cases the growth is considered to occur at a constant rate of volume increase and to be completed over the coarse time interval Δt’. **(Bottom Panel) Modelling intrinsic variability in inheritance:** Values of the parameters governing the kinetic transition constants (τ_I→J_) of the yeast growth and division steps are passed on from mother to daughter with a set degree of variability governed by three parameters (A_av_, σ_A_ and σ_B_) defined in Eqns. 3 and 4. The degree of variation is dictated by sampling from a Gaussian distribution the parameters which evolve in a lineage specific fashion. (**Upper**) **Schematic indicating how the potential for variation is modelled using a generalized parameter A:** A particular value from **A** is inherited in a particular lineage (red boxes) the value of which is shown using a grayscale distribution (A is sampled from a generation and lineage specific distribution **A**{A_av_(G),σ_A_(G, σ_B_)} (Eqn. 3 and 4)). (**Lower**) One example of a lineage and generation specific evolution of the distribution of potentialities **A**{A_av_(G),σ_A_(G, σ_B_)} after 10 generations with A_av_(G=1) = 0, σ_A_(G=1) = 0.01, σ_B_= 0.01.

Successful passage through these various life transitions (here generalized as I → J) is governed by a transition probability P(I→J) which is itself a function of time, t, and local yeast density, ρ_local_. To formulate the transition probability into algorithmic form we first decompose it into the product of two limiting probabilities (**Eqn. 2a**). The first limiting case involves the transition of an isolated yeast (zero local density) at some finite time t, defined in relation to the limiting minimum delay δ_I→J_. This transition probability is determined in a stochastic fashion on the basis of a first order process calculated using the characteristic time constant (**Eqn. 2b**) **[Hartwell and Unger, 1977; Lord and Wheals, 1980]**. The second limiting case considers the transition at infinite time^10^ but at non-zero local yeast density such that transition success is wholly determined by the ability of a growing yeast bud, or daughter cell, to overcome any virtual pressure generated by local yeast occupancy, due to either a preference for cohesion between yeast in a colony (so-called cell to cell contacts) or the formation of anchor points between the yeast and the plate^11^ [**Roy et al. 1991; Bony et al 1997**]. To model this virtual pressure aspect associated with yeast colony growth, all yeast obstructing either, the point of intended daughter formation or enlargement that could impede such growth, are first identified (**Fig. 2 – Middle Panel**). Knowledge of the number and placement of these surrounding yeast is then used to calculate a Metropolis-like weighted selection term based on a dimensionless^12^ energy, ΔE^*^, that factors in the requirement to push any (and all) obstructing yeast away from the point of daughter formation/enlargement (**Eqn. 2c**) **[Leach 2001]**. The energy term appearing in Eqn. 2c is calculated from the following parameters; ε - the reduced energy required to push one yeast segment a minimal distance 2R_D_ and min[N_(┴)_, N_(┬)_] - the smallest value from the set of the total number of obstructive yeast segments that have to be moved in one of the two opposing Cartesian directions (**Eqn. 2d**)^13^.

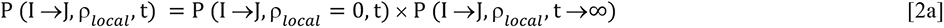

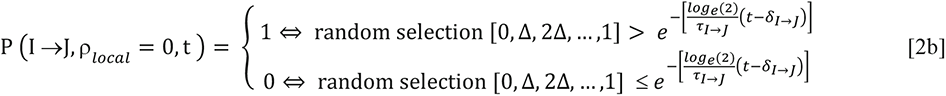

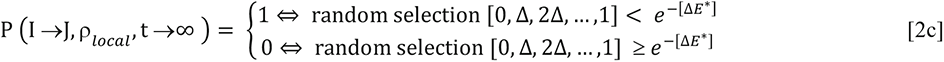

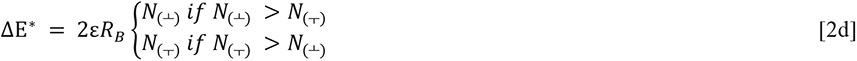

For the M → D transition the bud may appear on any one of six positions of the yeast faces that are perpendicular to the growth surface (**Fig. 2 - Middle Panel**). For the D → M transition, one of four potential positions, located a distance 2R_D_ from the center of the grown bud (and aligned parallel to the xy axes), is selected for extension to the mature asymmetric yeast form (**Fig. 2 - Middle Panel**). For all putative growth transitions, if P(I→J, ρ_local_ = 0, t) = 1, then all sites are tried via random selection without replacement until either P(I→J, ρ_local_, t®¥) = 1 or no further selections are available, in which case P(I→J,ρ_local_, t®¥) = 0 and the total probability in Eqn. 2a is set to zero i.e. P(I→J, ρ_local_, t) = 0.

#### Accounting for variability in yeast strain and individual cell characteristics

Typically, our simulations consider the growth of yeast that belong to a single strain type (with strain defined by degree of isogenic character / and morphology) [**Louis, 2016**]. At time zero, the yeast on the plate, N(t=0), is considered first generation, G = 1, and assigned a unique index, H = 1, associated with its time of appearance on the plate. For all subsequent divisions each indexed cell is assigned both a generation index, G = 2,3,.., a unique index H such that H ∈ [1,2, …, N(t)] and a specific cell lineage (written as a concatenated list of the particular indices of cells involved in the division chain). Together with the particular transitive state of the yeast (D, M) this information can be used to partially define the state and history for each yeast on the plate.

Despite their isogenic nature, individual members of the same yeast strain will exhibit variation in their growth and division patterns, due to differences in internal constitution (e.g. mutations, stochastic separation of cytosolic components during division) and the external micro-environment (e.g. temperature, plate medium, surrounding cell density etc.) **[Hartwell and Unger, 1977; Snijders and Pelkman, 2011; Cerulus et al. 2016; Mayhew et al. 2017]**. To include these variations within the model the transitive time constants are modified using a number randomly selected from a normal distribution **A{**A_av_(G), σ_A_(G)} characterized by a mean, A_av_, and a standard deviation, σ_A_, which are both in turn set as functions of the yeast generation index. At each stage of growth the individual yeast cell’s transition time constant is allowed to vary (**Eqn. 3**) (**Fig. 2 - Lower Panel**).

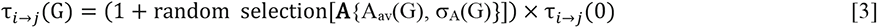

If A_av_(G) equals zero and σ_A_(G) = σ_A_(0) throughout the course of yeast colony formation then Eqn. 3 ensures limited variability in the individual time constants around their mean value τ_i→j_(0). However as different strains will undergo different extents of evolutionary drift (via genetic and epigenetic routes) through multiple rounds of cell division, the potential for changes occurring in both the mean and variance parameters is extant **[Cerulus et al. 2016; Mayhew et al. 2017]**. To include these variational characteristics within the model both the average and standard deviation, defining the normal distribution **A,** are allowed to vary through multiple rounds of cell division (**Fig. 2 - Lower Panel**) as per a recursive formula (**Eqn. 4**) which employs random selection from a second set of normally distributed numbers, **B**(0, σ_B_), defined by a zero mean and a standard deviation, σ_B_.

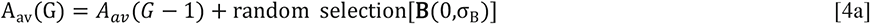

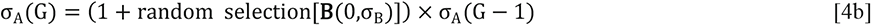

Inescapable tendencies towards greater entropy hardwired within the recursive identity of Eqn. 4 will lead to a broadening of the population level distribution of A_av_ and σ_A_ (as a function of generation time (G)).

#### Description of cell age in a chronological and a replicative sense

There are two definitions of age relevant to a discussion of cell growth. The first is chronological age, which describes the actual time for which the cell has been alive [**Bitterman et al. 2003; Longo et al. 2012;**]. Considerations of lineage can be important when the cell is exposed to any phenomenon which exhibits a time accumulated effect (e.g. time exposed to ultra violet light or chemical mutagen [**Longo and Fabrizio, 2011; Khokhlov, 2016**]). The second relevant measure of age is the replicative age, which describes how many times the cell has undergone the cell division process to produce a daughter cell [**Barton, 1950; Steinkraus et al. 2008]**. There are a number of interesting features about both of these measures of cell aging with regards to epigenetic phenomena. For instance one theory of cell division termed the maternal protective effect, is based on the observation of preferential retainment of damaged cellular components by the mother cell to allow the growing daughter cell to get off to the best possible start in life [**Kennedy et al. 1994; Steinkraus et al. 2008**]. Within MIL- CELL both linear and replicative ages are recorded for each cell in the virtual culture. Allowance can be made for the occurrence of cell death after set linear and replicative ages at which stage the yeast disappears from the culture plate. In the case where amyloid represents a disease of yeast both the growth parameters and the age at which the yeast cell dies can be modified in response to amyloid load [**Wickner et al. 2011; Douglas et al. 2011**].

#### Mesoscopic assignment index: Observable properties of yeast at the individual cell level

Taken all together Eqns 1-4 allow for the definition of an information set **Y** with each table row defining a particular yeast’s state (at the particle level of consideration) using the indexing scheme shown (**Eqn. 5**) (with the additional terms indicated by … to be added in a subsequent section).

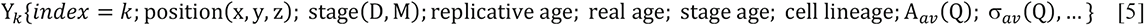

Up to this point we have described how to specify the physical location and division cycle particulars of each yeast cell within a growing colony. However, we have not yet specified any mechanism for describing information on the state of the amyloid prions existing within each yeast cell. To remedy this deficiency, in the next sections we describe a chemical rate model of amyloid prion growth and then explain how to integrate this modelling framework into the particle-level description of the yeast life cycle to generate a true multi-scale model that is able to inform on the amyloid prion population within the yeast during its growth and colony formation on a plate.

### (ii) Microscopic description of amyloid prion chemical processes

#### Chemical rate equations describing the behaviour of yeast prions within each yeast cell

Conforming to the original discoveries of Wickner, we assume that the fundamental basis of the transmissible prion unit in yeast is protein amyloid [**Wickner, 1994; King et al. 2004; Brachman et al. 2005**]. In keeping with this point, here we describe methods for simulating the nucleation and growth of amyloid from its monomeric protein precursor based on the numerical integration of a set of chemical rate equations (defined by such polymer nucleated/growth considerations) [**Hall and Edskes, 2012; Hirota et al. 2019**]. In what follows, we consider two general sets of amyloid growth equations that are capable of producing three distinctive prion kinetic behaviors observed in yeast described as ‘dissolution/munching’ (increased rate of endwise depolymerization), ‘inhibition of fiber breakage’ (reduced rate of internal fiber breakage) and ‘clumping’ (self-association of amyloid fibers) with this latter process associated with asymmetric segregation upon cellular division [**Zhao et al. 2018]** (**Fig. 3 - Upper Panel**). We describe in detail the elementary steps both common and particular to these three general behaviors before providing the relevant equation sets.

**Figure 3:**
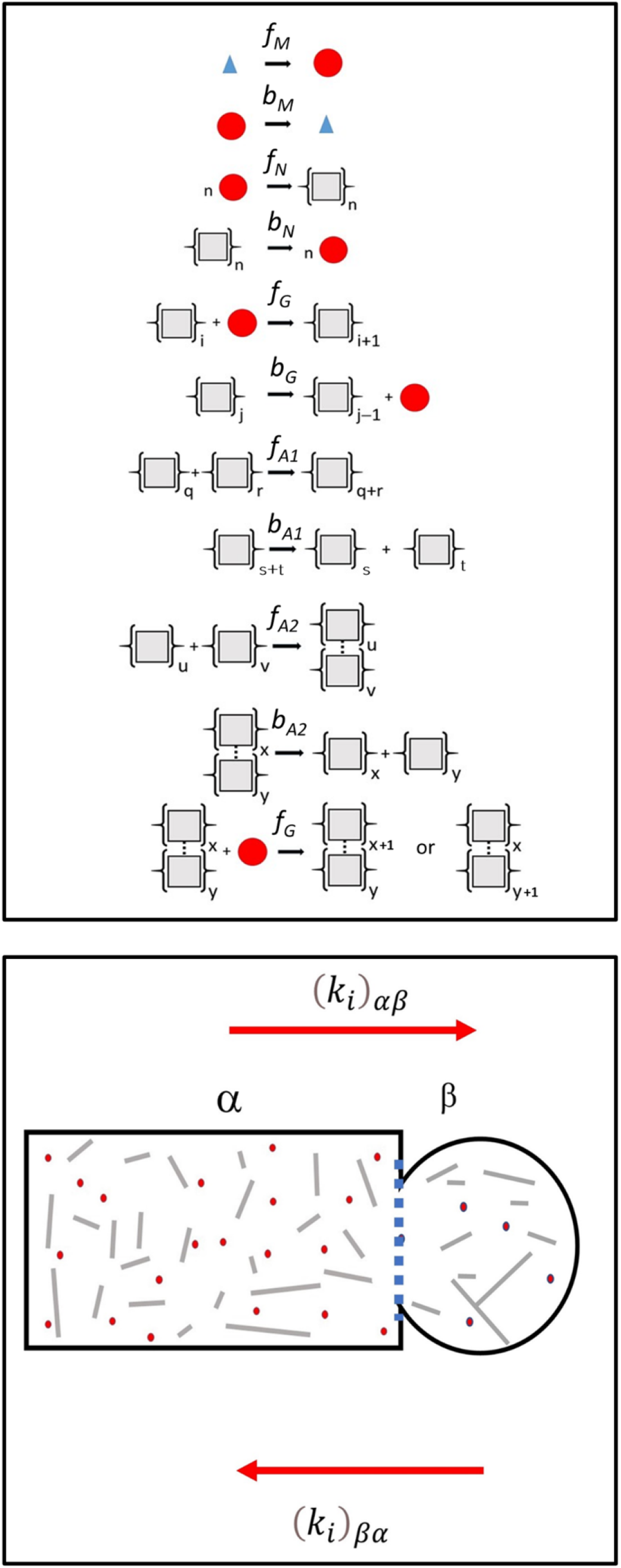
Kinetic models for amyloid formation and partition. **(Top Panel) Kinetic mechanism of amyloid formation within S. cerevisiae.** Amyloid formation is broken down into a series of forward and backward elementary steps respectively governed by rate constants f and b (with subscripts specific for the class of reaction). Under this governing mechanism, the formation of monomer (red circles) from amino acids (blue triangles) is governed by first order forward and backward constants, f_M_ and b_M_ (units s^−1^). Protein conversion to the amyloid structural state is designated by {grey squares}_j_ with the index indicating the number of monomers with the aggregate state. The formation of an amyloid structural nucleus, N, from n monomers (indicated by {grey square}_n_) is respectively governed by an n^th^ order forward constant f_N_ (units M^−(n−1)^s^−1^) and a first order backwards constant b_N_ (units s^−1^) (*in this work n is exclusively set to 2). Growth and shrinkage of an amyloid protofibril (single fiber) can occur either via monomer addition and monomer loss (with these steps respectively governed by a second order forward rate constant f_G_ (units M^−1^s^−1^) and a first order backward rate constant b_G_ (units s^−1^)) or via joining and scission of complete amyloid fragments (with these steps respectively governed by a second order forward rate constant f (units M^−1^s^−1^) and a first order backward of undergoing either breakage or monomer dissociation but can incorporate further monomer at a rate governed by the second order rate constant f_G_. Specifying zero and non-zero values for these individual steps allows for the user to specify different types of characteristic growth behavior comporting to certain classes of amyloid kinetics that can feature breakage, preferential endwise ‘munching/dissolution’ or amyloid fiber clumping [**Zhao et al. 2018**]. **(Lower Panel):** Modelling the component specific partition from the mother cell (termed the α phase) into the daughter cell (termed the β phase) and also from the daughter to the mother cell during the process of cell division. For each i^th^ class of chemical component i.e. i ∈ {M, N, A1, A2} a unique partition constant (having units of s^−1^) is specified in the α→β and β→α direction respectively as (k_i_)_αβ_ and (k_i_)_βα_ and the rate of migration of each class of component is determined using Eqn. 10 in conjunction with Eqn. 11.

*Protein production:* The formation and breakdown of protein monomer, M, from (and to) its constituent amino acids, is considered to be respectively regulated by first order rate constants, f_M_ and b_M_. The concentration of amino acids available for monomer production is itself parameterized in terms of the total mass concentration of amyloid, its basal set-point concentration (C_AA_)_basal_ at zero amyloid concentration and two empirical parameters, Ω and Ψ (**Eqn. 6**). This parameterization accounts for the fact that amyloid is believed harmful to the cell beyond a certain concentration [**Wickner et al. 2011; Douglas et al. 2011; McGlinchey et al. 2011**] therefore making its own production self-limiting at some extent of amyloid production^14^. We interpret this self-limiting aspect as a general decrease in metabolic function modelled as a decrease in the available pool of cell resources^15^.

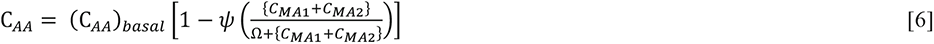

*Amyloid nucleation:* The initial amyloid nucleation process is considered to occur via an association event of molecularity n, governed by an n^th^-order^16^ association rate constant, f_N_. Nucleus dissociation is considered to be governed by a first-order dissociation rate constant, b_N_.

*Amyloid growth via monomer addition/loss:* Amyloid growth and dissociation is specified to occur in a simple manner via monomer addition to a single fibril end, governed by the second-order rate constant f_G_, and monomer dissociation from either of the fibril ends^17^, governed by a first-order rate constant b_G_.

*Amyloid growth (linear fiber addition/breakage):* Amyloid fiber growth and breakage may additionally occur via fiber end-to-end joining **[Binger et al. 2008]**, regulated by a second-order association rate constant, f_A1_, and internal breakage, governed by a first order site-breakage rate constant, b_A1_.

*Amyloid growth (lateral fiber addition/dissociation):* Amyloid ‘clumping’ may also occur by fiber lateral association with the forward reaction governed by a second order association rate constant, f_A2_, and the reverse reaction governed by the first order rate constant, b_A2_. The fiber clump is considered to be able to undergo growth via monomer addition to the two available ends (unidirectional growth) with this growth regulated by the second-order rate constant f_G_. Due to the potential additional stabilization of the fibers due to their lateral alignment, the clumped fibers are considered not to be able to undergo either monomer dissociation or fiber breakage in the stabilized clumped state.

*Species partition:* The monomer and amyloid components may migrate between mother and nascent daughter cells^18^ via differential partition during the budding phase resulting in daughter cell formation (**Fig. 3 - Lower Panel**). As such, each different chemical component was assigned two directional partition constants, (k_i_)_αβ_ and (k_i_)_βα_, (units of s^−1^) respectively referring to migration of component i from the mother (α phase) to the daughter (β phase) cell, and from the daughter to the mother cell. Due to the unequal volumes of the mother and growing bud a volume correction is applied to components entering and leaving the compartment with the larger volume (mother cell).

Towards formulating simplified mathematical representations of these events described in terms of their species concentrations, C_i_ and mechanistic [f_i_, b_i_] and partition, [(k_i_)_αβ_, (k_i_)_βα_] rate constants governing each set of forward and reverse elementary steps, we provide compartment specific (Q = α (mother) or β (daughter)), definitions for the amyloid number, {C_A1_}_Q_ and mass concentrations, {C_MA1_}_Q_ for a single filament, along with the equivalent concentration definitions for, the clumped (paired) filaments, {C_A2_}_Q_ and {C_MA2_}_Q_ (**Eqn. 7a-d**).

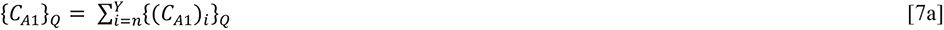

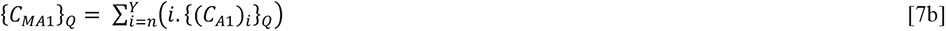

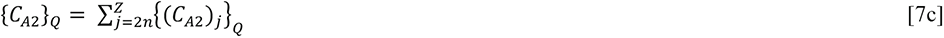

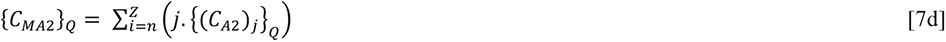

Using the species defined in Eqn. 7 along with a number of simplifying assumptions (which will be outlined below) we can write sets of rate equations that describe amyloid growth under relevant different limiting conditions [**Zhao et al. 2018; Greene et al. 2020]**.

#### Kinetic model capable of fiber scission and longitudinal/lateral fiber self-association (breakage and clumping)

A set of inter-related differential equations capable of describing amyloid fiber breakage, clumping and partition between the mother and daughter (with the partition term to be defined subsequently) can be derived upon making the following assumptions (i) that the nucleus size, n, is 2; (ii) the nucleus dissociation rate, b_N_, is equal to the dissociation rate of monomer from the fibril, b_G_ ; (iii) there is no positional dependence to the intrinsic breakage rate i.e. b_A1_ = b_G_ at all fracture points. These equations describe the phase specific rate of change in the concentration of monomer, (C_M_)_Q_ (**Eqn. 8a**), the number concentration of single filament (C_A1_)_Q_ (**Eqn. 8b**), the number concentration of paired filaments (C_A2_)_Q_ (**Eqn. 8c**), the mass concentration of single filaments (C_MA1_)_Q_ (**Eqn. 8d**) and the mass concentration of paired filaments (C_MA2_)_Q_ (**Eqn. 8e**) with the average size of the single and paired filaments (**Eqn. 8f** and **8g**).

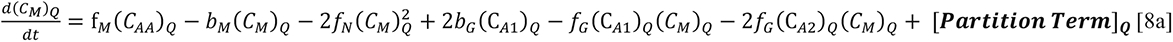

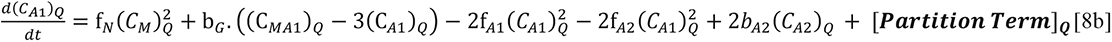

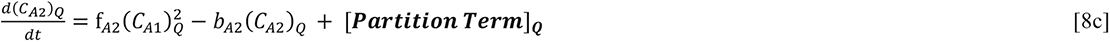

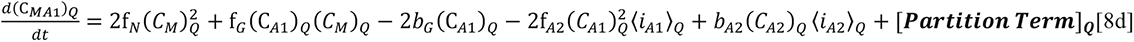

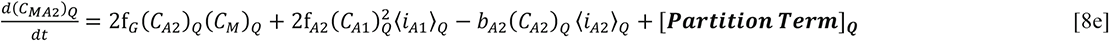

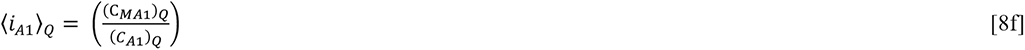

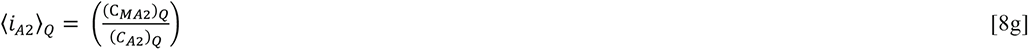

For the case of no fiber breakage {b_N_ = b_G_ = b_A1_ = 0 s^−1^} and no fiber joining or clumping {f_A1_ = f_A2_ = 0 M^−1^s^−1^} the chemical regime conforms to models of irreversible nucleated growth first pioneered by Oosawa and others [**Oosawa and Asakura, 1975**]. For the case where breakage is finite i.e. {b_N_ = b_G_ = b_A1_ > 0 s^−1^} without joining or clumping {f_A1_ = f_A2_ = 0 M^−1^s^−1^} the chemical regime conforms to a standard consideration of amyloid kinetics **[Hall and Edskes, 2009]**. When the joining and clumping rates take on a finite value {f_A1_ > 0 M^−1^s^−1^; f_A2_ > 0 M^−1^s^−1^} the fiber number concentration will be modified reflecting this fiber self-association **[Zhao et al. 2016; Hirota et al. 2019]**.

#### Kinetic model capable of position dependent breakage (endwise dissolution ‘munching’)

To simulate preferential endwise depolymerization of amyloid filaments (the so called endwise dissolution or ‘munching’ case postulated by Zhao et al. 2018) a previously developed mathematical model, able to account for position dependent differences in fiber fracture rate i.e. b_G_ ≠ b_A1_, was utilized [**Hall, 2020**]. Limited to the consideration of single filament growth this model is capable of describing the time dependence of monomer formation (**Eqn. 9a**), the number concentration of fibrils (**Eqn. 9b**) and the mass concentration of fibrils (**Eqn. 9c**) under the following assumptions (i) that the nucleus size, n, is 2; (ii) the nucleus dissociation rate, b_N_, is equal to the dissociation rate of monomer from the fibril, b_G_ ; (iii) the phase dependent concentration of the nucleus (C_2_)_Q_ is estimated by assuming an exponential shape of the fibril distribution (**Eqn. 9e**) for which the decay constant is calculated on the basis of knowledge of the average polymer degree (**Eqn. 9d,f**) **[Hall 2020]**.

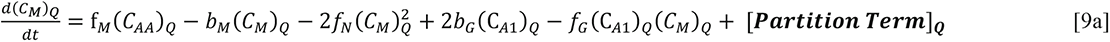

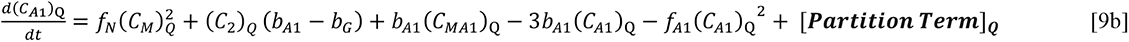

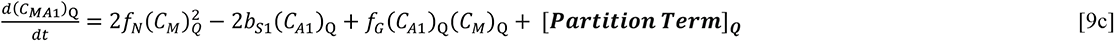

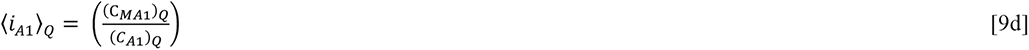

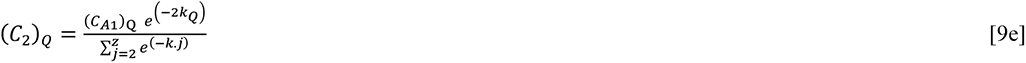

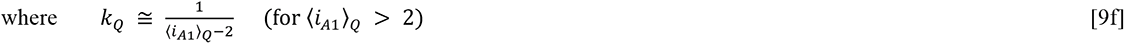

With the basic form of the kinetic equations for amyloid growth occurring in a fixed volume we now describe the functionalization of the partition term.

#### Partition of chemical components between the dividing mother and the daughter cell

Partition refers to the migration of the specified monomer and amyloid components between the mother and daughter cells during cell division **[Marchante et al. 2017; Zhao et al. 2018; Heydari et al. 2021; Greene et al. 2020]**. The partition terms shown in Eqn. sets 8 and 9 are necessarily different for each component and are also biased by a volume factor dependent on the volume compartment Q being discussed (either α (cell) or β (daughter) cell). The partition equation, written in terms of generalized parameters, is given (**Eqn. 10**)^19^

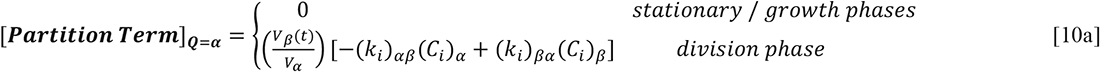

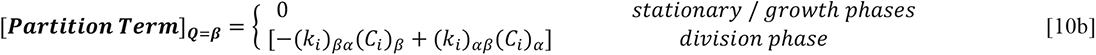

With suitable initial values, Eqn. sets 8 and 9 (with the partition terms described - **Eqn. 10**) are sufficient to describe characteristic growth patterns within yeast of unchanging physical dimensions (stationary phases) or alternatively a ‘frozen’ state of a dividing yeast. A multi-panel display (**Fig. 4**) describes the changes in the time dependent formation of monomer and amyloid as the kinetic parameters are systematically varied at fixed values of the other parameters in an unchanging solution vessel (e.g. as for a chemical beaker). However, in the form shown, Eqn. sets 8 and 9 cannot account for the kinetics of amyloid in a cell compartment undergoing volume change with time. To model this feature we need to recast the set of ordinary differential equations described by Eqn. sets 8 and 9 (in which the only independent variable is time), into a set of partial differential equations written in terms of the independent variables of time and volume. We begin by describing the total differential for the concentration of species k in terms of change in time and volume (**Eqn. 11a**). Approximating the derivative as a difference equation yields a functional form for the derivative (**Eqn. 11b**).

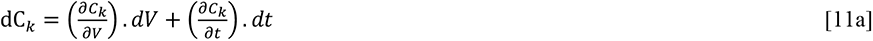

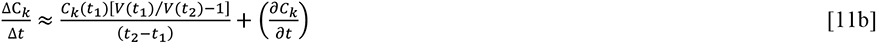

**Figure 4:**
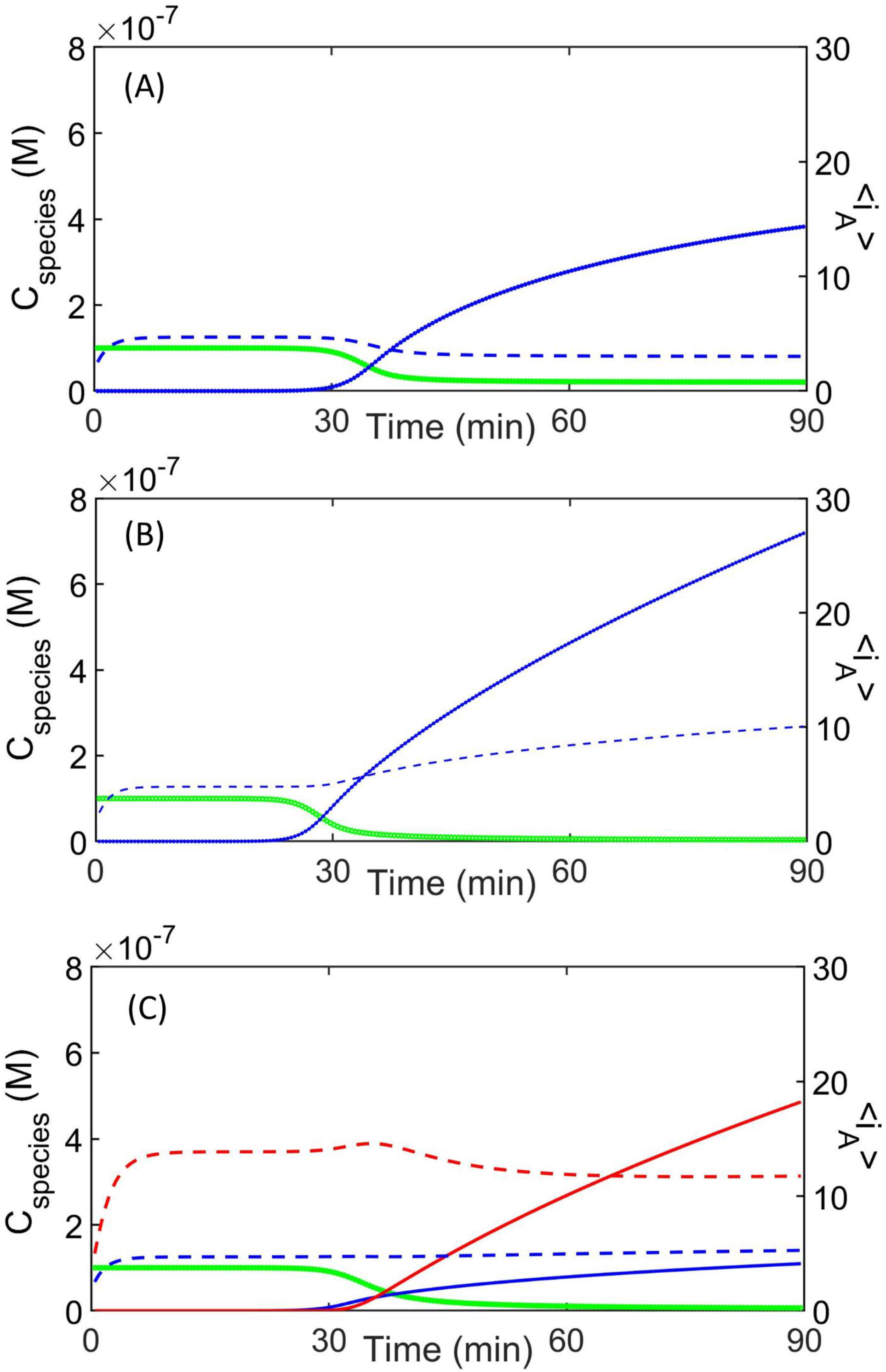
Modelling amyloid kinetics in a mother cell of fixed unchanging volume. Demonstration of how selection of mechanism and specification of parameters can determine the kinetics of amyloid growth within a single mother cell of unchanging dimensions over the average yeast lifetime (solid lines refer to concentration - green = C_M_, blue = C_MA1_ red = C_MA2_ ; dotted lines refer amyloid average size - blue = <i_A1_>, red = <i_A1_>). **(A) ‘Standard’ amyloid fiber breakage mechanism -** in which fibers can grow by monomer addition or fiber joining, and shrink by monomer loss and internal breakage [e.g. Hall and Edskes, 2009; Zhao et al. 2016] {in this particular example b_N_ = b_G_ = b_A1_ = 0.005s^−1^; f_G_ = 5×10^5^M^−1^s^−1^; f_A1_ = 0M^−1^s^−1^; f_A2_ = 0M^−1^s^−1^; b_A2_ = 0s^−1^ **(B) Differential ‘Munching’ of amyloid fibers** - in which the intrinsic rates of amyloid breakage are considered to occur differently at the end of the polymer and at internal sites [e.g. Hirota et al. 2019; Hall, 2020] {in this particular example internal breakage is considered greater than endwise depolymerization b_N_ = b_G_ = 0s^−1^; b_A1_ = 0.005s^−1^ ; f_G_ = f_A1_ rate constant b_A1_ (units s^−1^)). Growth and shrinkage of a clumped amyloid fiber of arbitrary degree x+y {indicated by a cross bridge existing between the grey squares of two amyloid single fibers of individual degree x and y) is set by second order forward, f_A2_ (units M^−1^s^−1^) and backward, b_A2_ (units s^−1^) rate constants. Clumped fibers are assumed to be incapable f_A1_ = 5×10^5^ M^−1^s^−1^; f_A2_ = 0M^−1^s^−1^; b_A2_ = 0s^−1^} **(C) ‘Clumping’ of amyloid fibers** - in which amyloid fibers can laterally align to form stabilized fibers that are incapable of breaking or undergoing monomer loss {in this particular example b_N_ = b_G_ = b_A1_ = b_A2_ = 0.005s^−1^ ; f_G_ = f_A1_ = f_A2_ = 5×10^5^ M^−1^s^−1^} **[Zhao et al. 2016; Hirota et al. 2019]**. Common simulation parameters f_M_ = 0.01s^−1^; b_M_ = 0.01s^−1^; f_N_ = 0.001M^−1^s^−1^; C_AA_ = 1×10^−7^M; C_M_ = 1×10^−7^M; ψ = 0.95; Ω = 1×10^−7^M.

Having described means for modelling the time evolution of the different amyloid components within each yeast cellular space we may supplement the yeast population information set **Y** (described in Eqn. 5) with additional time dependent chemical information, with the specific terms indicated (**Eqn. 12**).

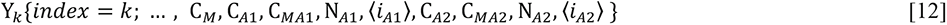

In Eqns. 1-12 we have described the yeast at the particle and the microscopic levels. In the next section we describe how these different levels of simulations are combined.

### (iii) Coupling of the mesoscopic and microscopic simulations

#### Algorithmic control of multiscale processes

Each life cycle associated cell volume change is considered to occur over the coarse time interval, Δt’, used in the particle model, with this volume increment broken into smaller time increments Δt determined by the number of time steps, N_steps_, used in the numerical integration routine for solving the amyloid kinetics (such that Δt = Δt’/ N_steps_. The particle model is updated at a regular interval Dt’. When the t + Dt’ time point reached, for each cell we carry out the following,

i. Apply a Metropolis-like test to determine the success or otherwise of the I → J transition (over Δt’)
ii. Solve the appropriate kinetic rate equations for amyloid aggregation within each yeast during the t → t + Δt’ period during which the I → J transition did (or did not) take place with either,

a. Eqn. 8 and 9 used for those periods of growth featuring fixed geometry, or
b. Eqn. 11 for periods of growth with changing geometry.
iii. For the case of a mother-daughter pairing the differential partition relation is used to account for uneven sharing of cellular contents at the time of septum closure (Eqn. 10).
iv. Next time advancement step in particle model (next Δt’)

The differential equations shown in Eqn. set 8-11 are evaluated using a modified-midpoint method in the numerical integration procedure [**Press et al. 2007**]. At the conclusion of each coarse interval Δt’ the end values are then used as the initial values in the next round of computation (upon reaching the next time interval).

#### Colour-based inference of the presence or absence of the yeast prion

In principle, yeast can display an observable phenotype that is caused by either the presence of prion (i.e. via a fluorescent screen in which the yeast are subsequently fixed and stained with a dye that is active upon encounter with amyloid [**Summers and Cyr, 2011]** or the amyloid forming protein is a fusion construct containing a fluorescent tag **[Zhao et al. 2018]**), or the absence of the monomer (in which the enzymatic activity is monitored [**Alberti et al. 2010**]). A number of biochemical assays can be applied/engineered within the yeast to make detection of the prion easy to carry out at the macroscopic level. Figure 5 describes the details of the assay for the identifying yeast cells containing the either the [PSI+] or [URE3] prion respectively made from the Sup35 and Ure2 proteins **[Schlumpberger et al. 2001; Alberti et al. 2010; Brachman et al. 2006**] (**Fig. 5a**). Broadly speaking, within an engineered yeast strain containing a premature stop codon within the ADE1 gene, the presence of the free monomer protein within the cytosol prevents expression of the Ade1p protein (coded by the ADE1 gene) which catalyses enzymatic breakdown of a coloured intermediate of the adenine biosynthetic pathway thereby causing the yeast cells to become red (also requiring that the growth media be supplemented with adenine). In the absence of free monomer (either Sup35 or Ure2p), such as when all monomer is in the inactive amyloid form, the dye converting enzyme is made and the yeast cells turn white (also meaning that the prion containing yeast can grow on adenine free media).

**Figure 5:**
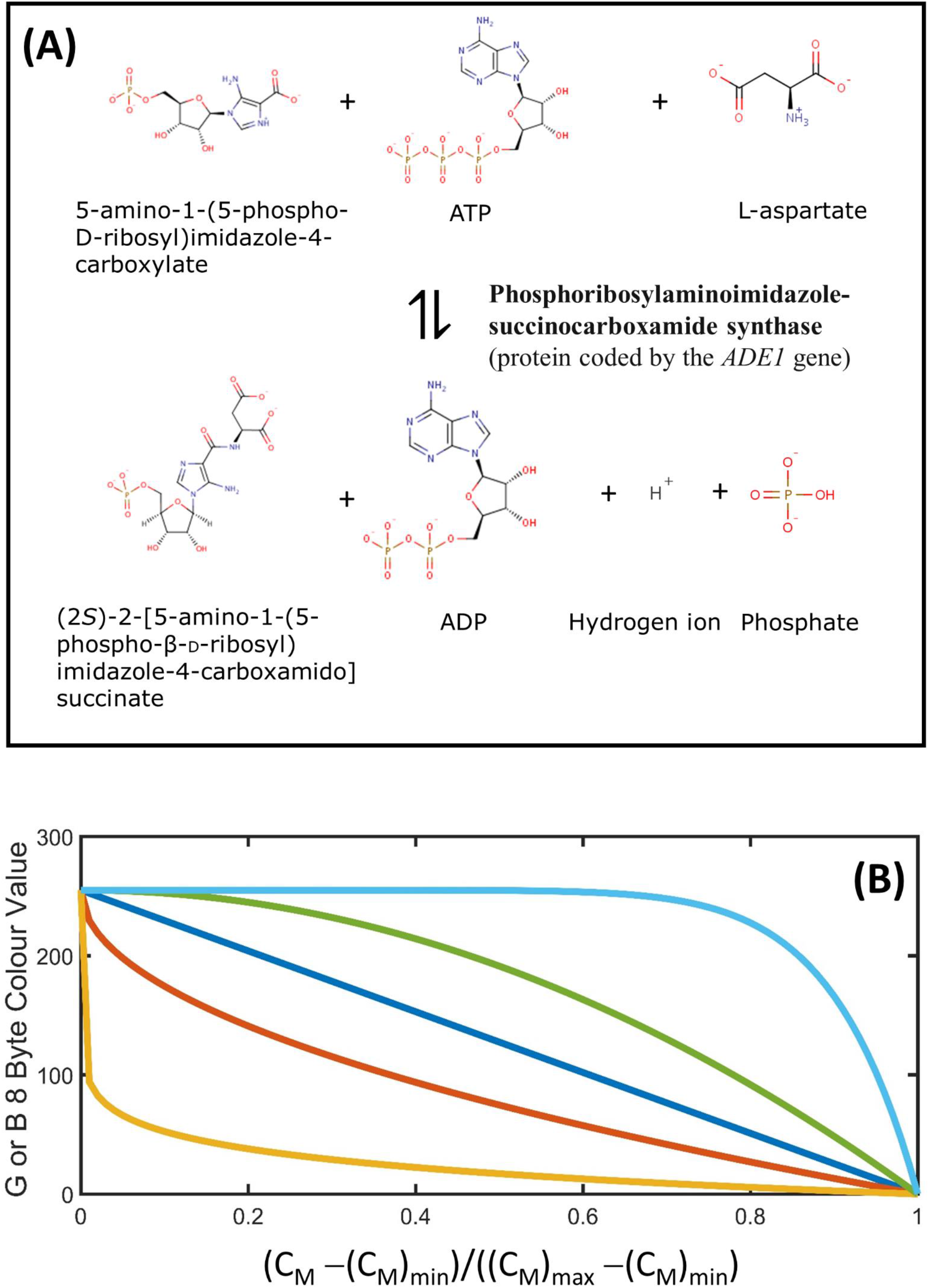
The basis of the color development reaction used for both the [PSI+] and [URE3] prion red/white colony screening assay in mutant yeast containing either a premature stop codon within the ade-1 gene [Alberti et al. 2010] or control of the ade-1 gene placed under a Gln3p promoter that is negatively regulated by the Ure2 protein [Brachman et al. 2006]. **(A)** In the case of the [PSI+] assay the translation complex bound to an ADE1 mutant mRNA containing a premature stop codon is stalled upon binding of soluble Sup35 thereby preventing expression of the functional ADE1 gene product (the phosphoribosylaminoimidazole-succinocarboxamide synthase protein also known as Ade1p). In the case of the [URE3] assay either the ADE1 is placed under the control of the Gln3/ DAL5 promoter. Ure2p binds to the Gln3 transcription factor and prevents it from entering the nucleus thereby preventing translation and expression of the ADE1 gene product. When present this protein catalyses the conversion of 5-amino-1-(5- phospho-D-ribosyl)imidazole-4-carboxylate to (2S)- 2-[5-amino-1-(5-phospho-β-D-ribosyl) imidazole-4- carboxamido] succinate with the former causing the yeast to be red in color. The presence of a prion within the yeast cytosol acts to sequester soluble monomer into the amyloid (non-functional state) and allows for enzymatic color removal by the ade-1 coded enzyme. This causes the yeast to take on a white color. **(B)** Modelling the relationship between the soluble monomer concentration and the degree of color development. Many scientists relate the color of the yeast colonies to the strength of the prion phenotype with color gradings such as strong (white), weak (pink) and absent (red). To provide a quantitative relation we institute a gamma function [Eqn. 13] that relates observed color to the cytosolic soluble monomer concentration between a minimum and maximum limit. Values of γ can vary between the limits (0,∞) providing a potential nonlinear dependence between observed color and monomer concentration. Lines shown represent γ = 0.1 (orange), γ = 0.5 (brown), γ = 1 (blue), γ = 2 (green), γ = 10 (aqua).

To quantitatively model the colour assay we have utilized a simple power dependence of the fractional level of change in monomer between minimum and maximum threshold limits (Eqn. 13). In signal processing such an equation is termed the gamma transformation with values of γ =1 denoting a linear dependence, γ > 1 denoting a ‘cooperative’ non-linear dependence and values of γ < 1 denoting an ‘anti-cooperative’ non-linear dependence **[Poynton 1998]**. A unique mapping is assigned in colour space by relating the fractional transition in monomer to the fractional transition in white to red colour space based on an 8-byte RGB representation with white denoted as [255,255,255] and red as [255,0,0] (**Fig. 5b**).

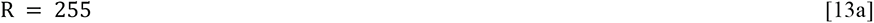

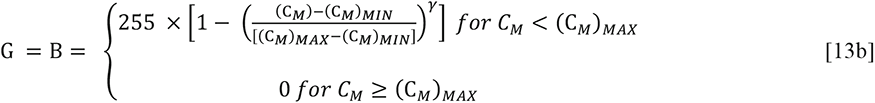

The color transform described in Eqn. 13 allows for a simple and direct means for simulating the amyloid status of any particular yeast via ‘visual’ inspection of the colony [**Brachman et al. 2006; Alberti et al. 2011**]. In what follows we will utilize both the particle level colour assessment and the microscopic chemical description to examine the epigenetic consequences associated with different relative parameter regimes of yeast and amyloid growth.

#### What can the MIL-CELL computational tool do?

Having explained the physical basis of the MIL-CELL model we now describe some of the types of virtual experiments and analysis results achievable with this software tool. Although not limited to the following we describe four types of potentially interesting in silico experiment achievable within MIL-CELL that conform to (i) Colony interrogation, (ii) Confluence analysis, (iii) Lineage and fate mapping, and (iv) Yeast curing experiments.

##### (i)#Colony interrogation

After specifying the yeast properties and initial conditions yeast cells are grown virtually, allowing for the spread of amyloid prions within them. A useful aspect of simulation is that it provides near complete knowledge of the behaviour of the system in a manner that is frequently not achievable even with the most carefully designed experiment. Taking advantage of this aspect of the MIL-CELL computational tool we have implemented a relational database that allows the user to investigate the properties of the yeast colony according to multiple parameters that describe properties of the cell (e.g. number of cells of a particular generation, their inherited variability, time of birth, time as daughter and time as mother) and all aspects relating to their cell contents (e.g. concentration and distributive state of the monomer and various amyloid forms) (**Fig. 6**).

**Figure 6:**
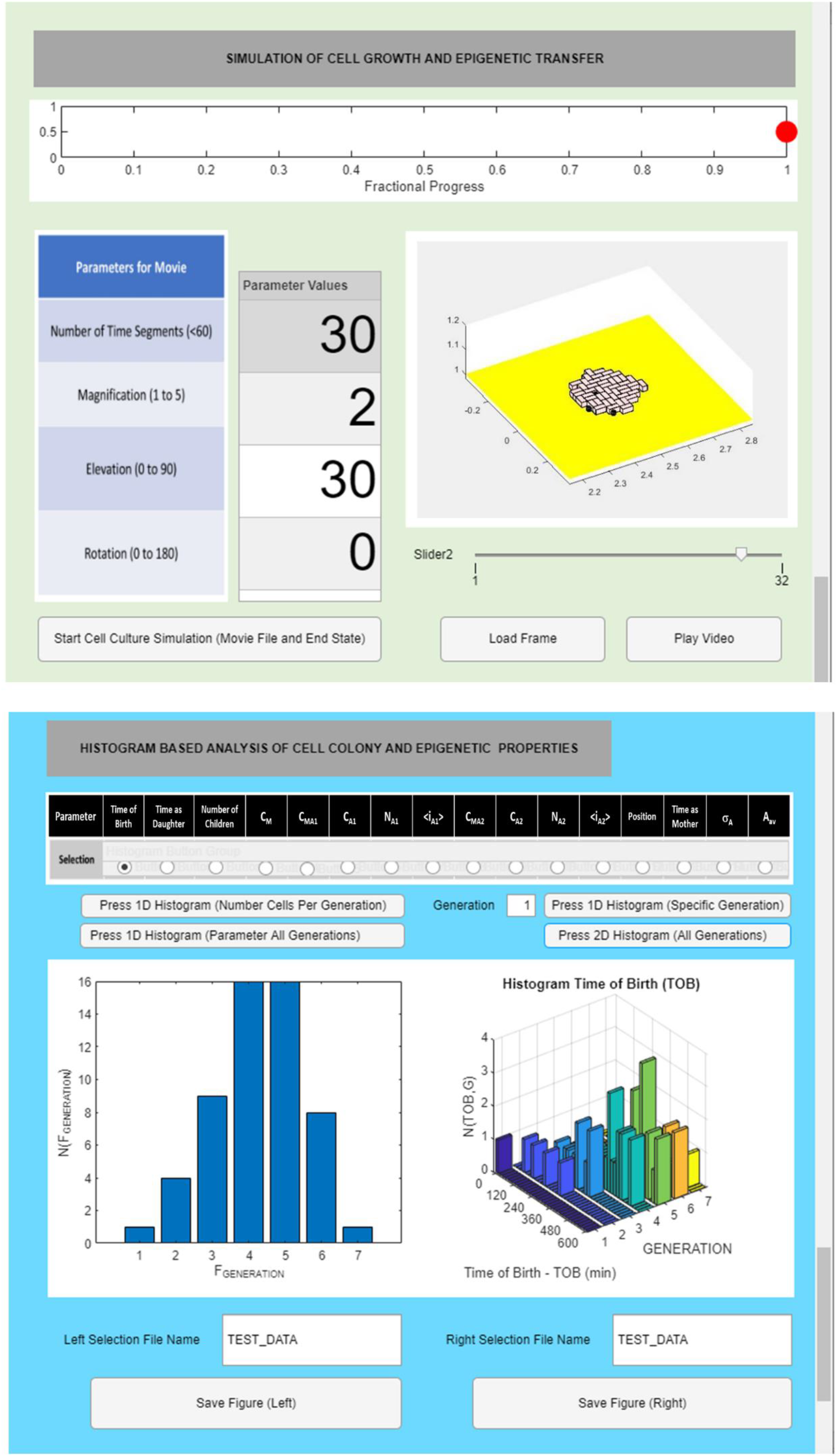
Colony interrogation using the MIL-CELL computational tool. **(TOP)** Cell culture experiment is simulated by specifying the number of coarse time increments (Δt’ = 20 minutes). A movie of the cell growth pattern is automatically generated and the various slides of this growth pattern can be examined frame by frame in the display window. **(BOTTOM)** A range of one- and two-dimensional histograms can be created based on cell generation and up to sixteen other selectable properties of yeast cell population. Selectable properties include (i) Time of birth; (ii) Time as daughter; (iii) Number of children; (iv) C_M_; (v) C_MA1_; (vi) C_A1_; (vii) N_A1_; (viii) <i_A1_>; (ix) C_MA2_; (x) C_A2_; (xi) N_A2_; (xii) <i_A2_>; (xiii) Position; (xiv) Time as mother; (xv) σ_A_; (xvi) A_AV_

##### (ii)#Confluence analysis

Depending on the degree of interaction of the growing cells with the support medium and/or other nearby cells there will be some difference in preference for growth occurring external to the colony (i.e. at the edges) vs. internally (thereby requiring displacement of surrounding cells). By varying the confluence parameter ε (shown in Eqn. 2d) we can alter the growth patterns of the yeast to either respect the local confluence condition^20^ or not respect local confluence and thereby grow in a less controlled fashion without effect from their surrounding neighbouring cells [**Verstrepen and Klis, 2006**] (**Fig. 7**).

**Figure 7:**
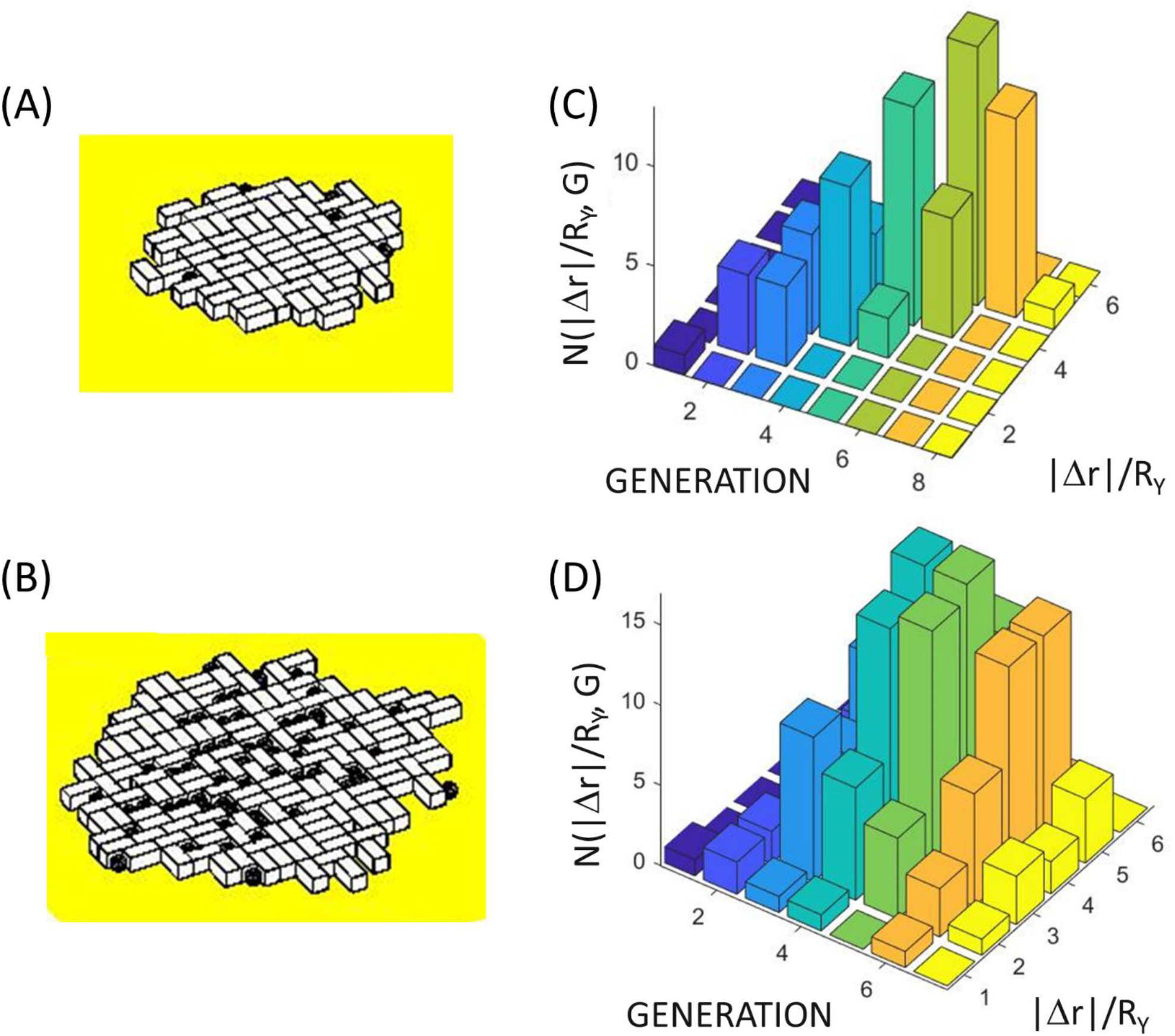
Examining the effect of the confluence parameter, ε, on the position and number of cells grown within a yeast colony over a period of 700 minutes. **(Left Panels)** Result of yeast cell colony growth for the case of **(A)** confluent growth conditions (ε = 1×10^12^ m^−1)^ and **(B)** non-confluent growth conditions (ε = 1×10^5^ m^−1^ ) (see Eqn. 2d). Note loss of confluency results in a faster rate of growth i.e. more cells produced. **(Right Panels)** Analysis of the position of yeast growth within the colony shown as a histogram of yeast generation and absolute displacement from the weighted center of the yeast colony normalized by the radius of the yeast, |Δr|/R_Y_). Panels display the cases of **(C)** confluent growth (analysis of simulation described in (A) with ε = 1×10^12^ m^−1^), and **(D)** non-confluent growth (analysis of simulation described in (B) with ε = 1×10^5^ m^−1^). Note confluent growth results in new generations being produced at edges of the colony whereas loss of confluency results in new generations being produced both within and at edges of the colony. [**Common yeast growth parameters [τ_D→M_ = 20 min, δ _D→M_= 0 min; τ_M→D_ = 20 min, δ _M→D_= 50 min**]

##### (iii)#Cell lineage and fate mapping

When conducting epigenetic linkage analysis a defined history of cell lineage is crucial for understanding how the epigenetic properties of a particular cell were determined by its immediate ancestors. In both the analysis of simple cell culture experiments and more complex pathways of growth and division in multicellular organisms such a history is known as a cell lineage map **[Woodworth et al. 2017]**. Within the MIL-CELL program a particular cell within the two-dimensional cell culture experiment can be interrogated by button click to identify the unique index k (**Eqn. 5**) assigned to it in order of its birth (**Fig. 8a**). Upon selection of the ‘Complete Lineage’ option a history of the chosen cell (that traces back each mother daughter pairing to the original cell (k=1)) will be shown in a new window (**Fig. 8b**). This graph can be used to identify the time of birth of each ancestor and the time course of the epigenetic components (chemical contents in terms of monomer, amyloid protofilaments and clumped amyloid fibers) as they are transferred through the lineage. A second type of forward-looking cell analytical tool, known as a cell fate map is also available within MIL-CELL (**Fig. 8c**). Based on the description of all the offspring produced by a particular cell, the forward-looking fate map can reveal differences in inherited epigenetic components between sibling cells due to birth-order effects **[Cerulus et al. 2016; Mayhew et al. 2017**].

**Figure 8:**
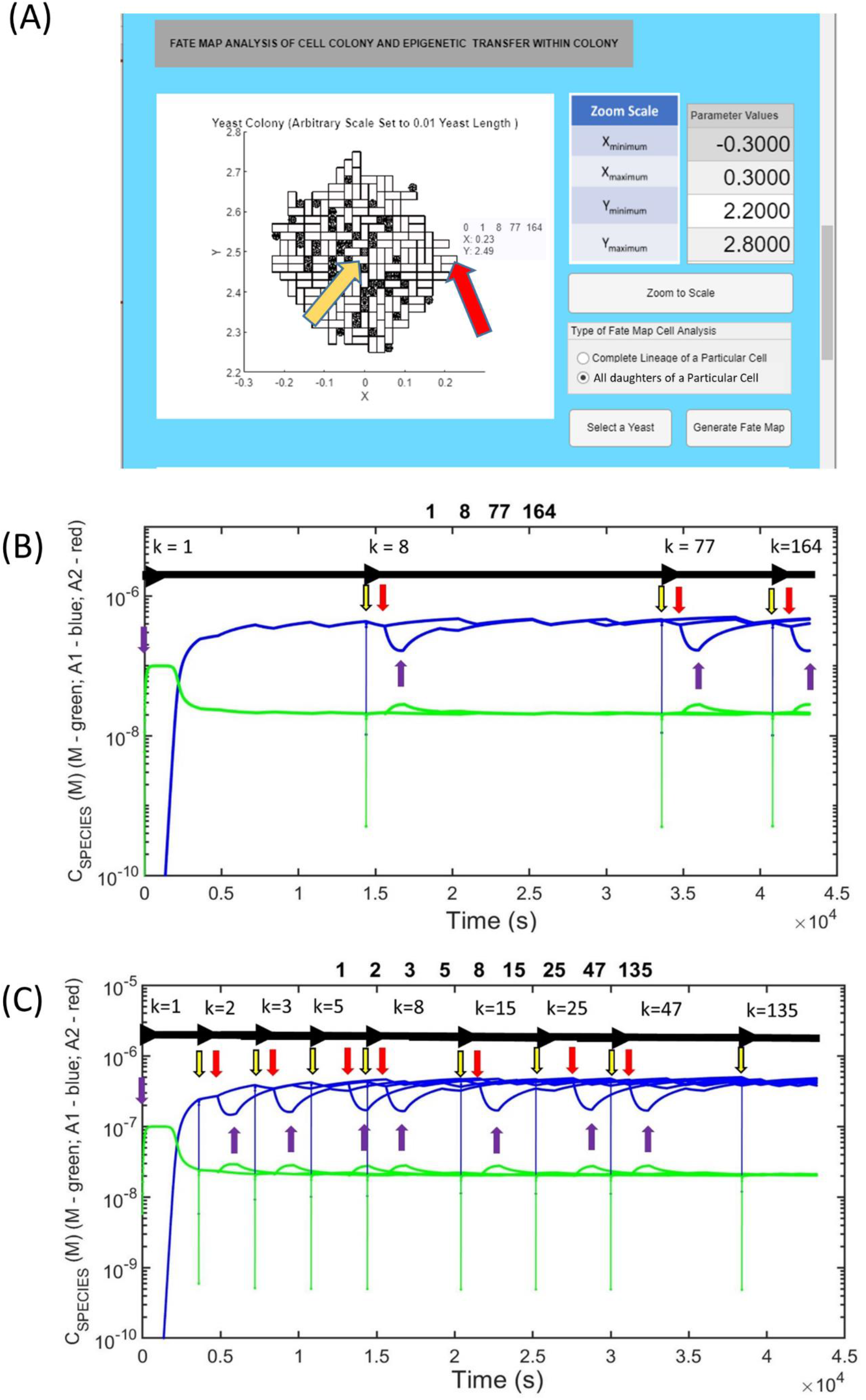
Examining the history and future of a cell via lineage and fate mapping: **(A) Virtual cell culture can be interrogated** - Top view of yeast cell colony growth (parameters given below). Red arrow indicates a selected cell k = 164 with lineage [1,8,77,164] (meaning that k = 164’s mother was k = 77, its grandmother was k = 8 and great grandmother was k = 1). The yellow arrow reveals cell k = 1. **(B) Cell lineage map:** A backward looking cell lineage, describing both the timing of the cells appearance and the concentration of amyloid protofilament and monomer concentration at each stage, can be constructed. As an example we show the lineage of cell k = 164. Horizontal black arrows indicate start of generation. Vertical arrows: Yellow - Initial point of budding t growth of daughter; Red – Start of growth of daughter cell to form mature cell (capable of becoming a mother); Purple - Formation of a fully grown mature cell. Note the spike in concentrations at the point of budding (nascent daughter cell) as amyloid and monomer contents partition into the bud from the mother cell. Note also the decrease in concentration of amyloid species and relative increase in monomer concentration during the increase in volume of the yeast cell during the daughter to mother transition. Following this the reverse behavior is seen as the concentration of amyloid re- establishes a pseudo-equilibrium in cells of unchanging volume. **(C) Cell fate map:** For a particular cell a forward- looking cell fate map can be constructed that describes the timing of the birth of all daughters, and the nature of the transfer of the cell’s internal contents (amyloid, monomers etc.). As an example we show the cell fate map for cell k = 1; Horizontal and vertical arrows are as for (B) with the exception that all sister cells are the same generation. Note for the present case a steady state level of amyloid is reached. **Cells grown for 700 minutes; Common yeast growth parameters [τ_D→M_ = 20 min, δ _D→M_= 0 min; τ = 20 min, δ = 50 min]; Kinetic parameters – Mechanism set at standard breakage [**f_N_ = 0.001M^−1^s^−1^; b_N_ = b_G_ = b_A1_ = 0.005^s−1^; f_G_ = 5x10_5_M^−1^s^−1^; f_A1_ = 0M^−1s−1^; f_A2_ = 0M^−1^s^−1^; b_A2_ = 0s^−1^ ; C_AA_ = 1x10^−7^M; C_M_ = 0M; ψ = 0.95; Ω = 1x10^−7^M; f_M_ = 0.01s^−1^; b_M_ = 0.01s^−1^]; **Cell partition parameters** – (k_i_)_αβ_ = (k_i_)_βα_ = 1 s^−1^ for all diffusible components. **Cell partition parameters** – (k_i_)_αβ_ = (k_i_)_βα_ = 1 s^−1^ for all diffusible components. **Cell variation parameters** [A_av_(G=1) = 0, σ_A_(G=1) = 0.01, B_av_ = 0 σ_B_= 0.01]. **Cell confluence parameter** [ε = 1×10^5^ m^−1^].

##### (iv)#Yeast curing experiments

A type of in yeast dilution experiment, known as yeast curing, was used to establish both the cytosolic location, and the amyloid prion nature of the epigenetic phenomenon **[Eaglestone et al. 2000; Wegrzyn et al. 2001; Byrne et al. 2009; Greene et al. 2020]**. In this experiment yeast are allowed to divide a number of times, with each generation tested for the presence of the epigenetic trait. The general hypothesis of the yeast curing experiment is that with each division cycle the contents of the cytosol are shared between an increased volume of cytoplasm (that of both the mother and newly formed daughter cells) and therefore undergo dilution [**Eaglestone et al. 2000; Cole et al. 2004; Byrne et al. 2007; Byrne et al. 2009**]. In practice, this experiment can be carried out either by growing several generations in solution with intermittent analysis via plating or via direct analysis of the yeast phenotype as it divides on the plate (known as a colony splitting/sectoring experiment) [**Sharma and Liebman, 2012**]^21^. The MIL-CELL program can simulate both forms of the yeast curing experiment with three examples of complete to partial curing generated by different mechanisms shown as **Fig. 9a-c**. A different manner for visualizing the yeast curing experiment involves plotting the fraction of cells not exhibiting the epigenetic phenotype against their generation number **[Eaglestone et al. 2000; Sharma and Liebman, 2012]**. To facilitate this representation MIL- CELL allows the user to specify the nature of the epigenetic marker (e.g. concentration of monomer or alternatively the concentration or number of prions) and to decide on the value of the binary classifier (i.e. what value demarcates the binary evaluation ([PSI+] or [psi−] of the epigenetic characteristic) (**Fig. 10**). Different specification of the type and value of this binary classifier can drastically affect the nature of the yeast curing curve.

**Figure 9:**
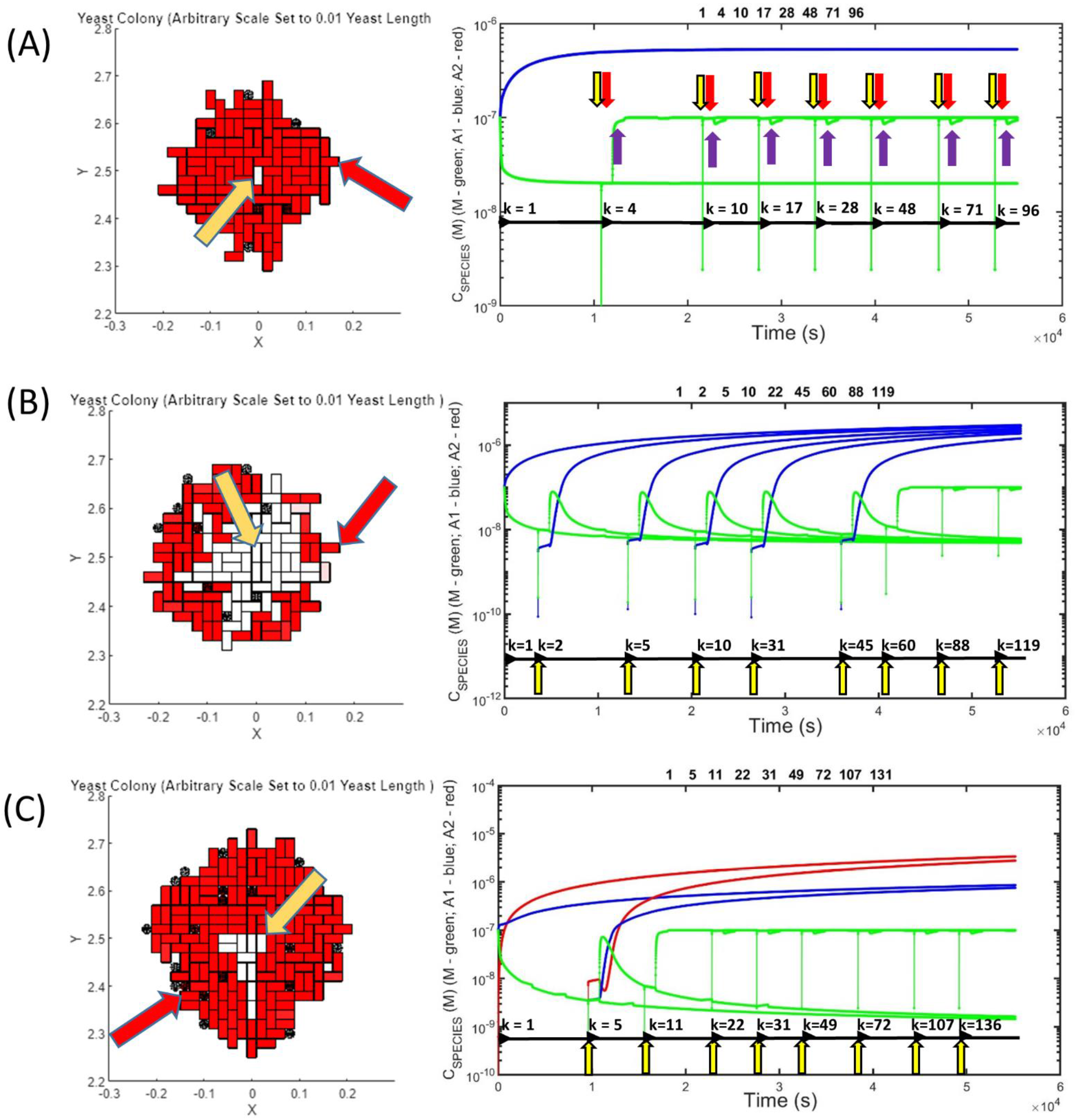
Examples of yeast prion curing experiment capable of being simulated in MIL- CELL. **(A) Yeast prion curing due to failure of amyloid partition from mother to daughter:** In this case the yeast lacks the facility for transmitting amyloid via partition from the mother to daughter cell. Left Hand Side (LHS) figure describes the simulated 2D-cell culture. The original amyloid prion containing cell (k=1) is shown existing in the center of the culture (orange arrow). A particular cell (k=96) chosen for the lineage mapping is shown at the periphery (red arrow). Right Hand Side (RHS) figure shows the lineage map between cells 1 and 96. The vertical yellow purple and red arrows respectively indicate the starting point of cell division, starting point of the daughter to mother transition and the completion of the daughter to mother transition. The horizontal black arrows describe the start of a new cell (from the formation of the bud with the k index provided immediately). **Unique parameters – Mechanism set to ‘standard breakage’ [**f_N_ = 0.001 M^−1^s^−1^; b_N_ = b_G_ = b_A1_ = 0.005 s^−1^; f_G_ = 5×10^5^ M^−1^s^−1^; f_A1_ = 0 M^−1^s^−1^; f_A2_ = 0M^−1^s^−1^; b_A2_ = 0s^−1^]; **Cell partition parameters** – (k_A1_)_αβ_ = 0 _s−1._ **(B) Partial yeast prion curing due to limited amyloid partition:** A finite (but limited) partition of amyloid from mother to daughter, (k_A1_)_αβ_ = 0.001 s^−1^, results in partial yeast curing and colony sectoring. LHS figure shows virtual yeast colony with orange and red arrows respectively describing cells k=1 and k= 119. RHS figure shows lineage map between cells k=1 and k=119. Note that the loss of the prion occurs at the k=45→k=60 cell division process. Vertical and horizontal arrows as per Fig. A. **Unique parameters – Mechanism set to ‘standard breakage’ [**f_N_ = 0.001 M^−1^s^−1^; b_N_ = b_G_ = b_A1_ = 0.005 s^−1^; f_G_ = 5×10^5^ M^−1^s^−1^; f_A1_ = 0 M^−1^s^−1^; f_A2_ = 0M^−1^s^−1^]; **parameters** – (k_A1_)_αβ_ = 0.0065 s^−1^. **(C) Partial yeast prion curing due to amyloid clumping:** The absolute discrete particle number of yeast prions can be decreased by lateral association of single amyloid fibrils to form ‘clumped’ fibers. A lower absolute number of fibers will decrease the transmission likelihood of prions during cell division. In this case such amyloid clumping results in colony spotting – the existence of a limited white coloured region exiting within a larger red background [**REFs**]. LHS figure virtual yeast prion curing experiment, orange and red arrows respectively describing cells k=1 and k= 131. RHS figure shows lineage map between cells k=1 and k=131. Prion loss occurs at the k=5→k=11 cell division process. Vertical and horizontal arrows as per Fig. A. **Unique parameters – Mechanism set to ‘clumping’ [**f_N_ = 0.001 M^−1^s^−1^; b_N_ = b_G_ = b_A1_ = 0.005 s^−1^; f_G_ = 5×10^5^ M^−1^s^−1^; f_A1_ = 0 M^−1^s^−1^; f_A2_ = 0M^−1^s^−1^; f_A2_ = 5×10^4^ M^−1^s^−1^ M^−1^s^−1^; b_A2_ = 0.001s^−1^]; **Cell partition parameters** – (k_A1_)_αβ_ = 0.0065 s^−1^. **Common yeast growth parameters [τ_D→M_ = 20 min, δ _D→M_= 0 min; τ_M→D_ = 20 min, δ _M→D_= 50 min]; Common kinetic parameters** C_AA_ = 1×10^−7^M; C_M_ = 0M; ψ = 0.95; Ω = 1×10^−7^M; f_M_ = 0.01s^−1^; b_M_ = 0.01s^−1^]; **Common partition parameters** – (k_i_)_αβ_ = (k_i_)_βα_ = 1 s^−1^ for all diffusible components unless otherwise specified. **Cell variation parameters** [A_av_(G=1) = 0, σ_A_(G=1) = 0.01, B_av_ = 0 σ_B_= 0.01]. **Cell confluence parameter** [ε = 1×10^12^ m^−1^]. **Cells grown for 900 minutes.**

**Figure 10:**
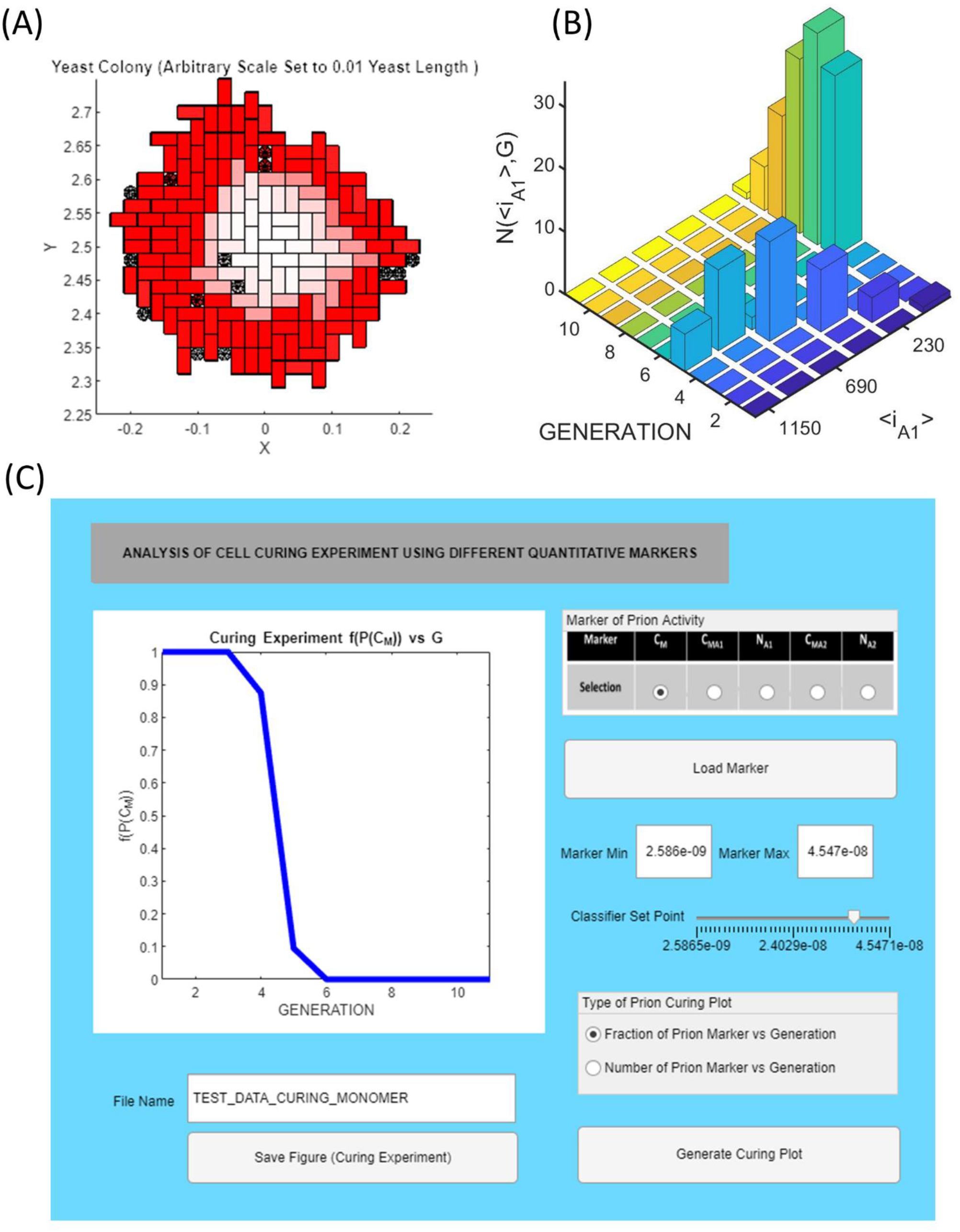
Description of the yeast prion curing experiment via a curing curve in MIL-CELL: The yeast prion curing experiment is typically analysed using a curing curve in which the fraction of cells exhibiting the epigenetic marker are plotted against their generation number. **(A) Yeast prion curing reflecting decrease in Hsp104 function instigated by inclusion of Guanidine HCl (GuHCl) in growth medium:** Inclusion of GuHCl leads to down regulation of the function of Hsp104 an active chaperone protein responsible for cutting yeast prion amyloid fibers into smaller pieces [Wegrzyn et al. 2004; Byrne et al. 2007]. The inability to fragment the amyloid fibers leads to radial dilution (in the case of confluent growth) of the prion amyloids with subsequent generation number. Yeast growth parameters [τ_D→M_ = 20 min, δ _D→M_= 0 min; τ_M→D_ = 20 min, δ _M→D_= 50 min]; Common kinetic parameters: C_A1_ = 1 ×10^−7^M; <i_A1_> = 5; C_AA_ = 1×10^−7^M; C_M_ = 0M; ψ = 0.95; Ω = 1×10^−7^M; f_N_ = 0 M^−1^s^−1^; b_N_ = b_G_ = b_A1_ = 0 s^−1^; f_G_ = 5×10^5^ M^−1^s^−1^; f_A1_ = 0 M^−1^s^−1^; f_A2_ = 0M^−1^s^−1^; b_A2_ = 0s^−1^; f_M_ = 0.01s^−1^; b_M_ = 0.01s^−1^]; Partition parameters – (k_i_)_αβ_ = (k_i_)_βα_ = 1 s^−1^ for all diffusible components. Cell variation parameters [A_av_(G=1) = 0, σ_A_(G=1) = 0.01, B_av_ = 0 σ_B_= 0.01]. Cell confluence parameter [ε = 1×10^12^ m^−1^]. Cells grown for 900 minutes. **(B) Description of the size of the amyloid as a function of yeast generation number:** A two-dimensional histogram of the number of cells of a certain generation possessing amyloid of a certain relative size <i_A1_> (in relation to monomer). We note that due to the chosen parameters reflecting GuHCl induced curing the yeas prions undergo dilution whilst also increasing in average size (i.e. due to the fact that b_N_ = b_G_ = b_A1_ = 0 s^−1^ yet f_G_ = 5×10^5^ M^−1^s^−1^). **(C) Screenshot of the MIL-CELL curing curve program section:** MIL-CELL offers a choice of five different markers for generation of the curing curve, C_M_, C_MA1_, N_A1_, C_MA2_, N_A2_, which can be presented in either fractional or cell number format. MIL-CELL features an option known as a ‘binary classifier’ which allows the user to decide what value of the marker determines a cured vs. non cured yeast.

The four just-described examples provide some insight into the potential usefulness of the MIL-CELL program for modelling various types of yeast prion experiments. In the next section we discuss how MIL-CELL compares with other models of yeast prion growth and transmission and how it may be applied more generally to other biological and disease phenomenon.

## Discussion

The starting intention of the present work was to describe how the MIL-CELL software could be used to model epigenetic effects mediated by the transmission of amyloid prions within yeast. However, due to the generality of the MIL-CELL modelling approach it has not escaped the authors’ notice that MIL-CELL may have a, not insignificant potential, to provide insight into a diverse array of general phenomena associated with eukaryotic cell growth and asymmetric division of cell contents in a manner that lies beyond the present discussion just associated with yeast prions. To place our work in both specific and wider contexts we have approached this discussion in the following manner. We first discuss MIL-CELL features in relation to the large numbers of models of yeast growth and division up to, and including, very recent models which feature prion growth and transfer. After presenting the strengths and weaknesses of the MIL-CELL program in relation to those other approaches we then place our focus on the biology of the processes modelled and compare our reduced description to current best understanding of how these complex processes occur in actuality. Finally, we discuss the potential of MIL-CELL in a wider context, by speculating on how its current (and future) capabilities might provide insight into a number of basic biological phenomena (such as cellular differentiation and cellular variability within a population) as well as cellular processes associated with disease (such as amyloidosis, cancer and mitochondrial dysfunction).

### (i)#MIL-CELL as a tool for modelling yeast growth/division and prion growth/transmission

To the best of our knowledge, MIL-CELL is the only model in existence that explicitly describes both (i) the spatial relationship between each yeast as they grow and divide in culture, and (ii) the time dependent chemical kinetics of amyloid prion growth and transmission within and between yeast. To achieve this feat MIL-CELL employs a multi-scale approach, meaning that it basically comprises two models in one, and as such we discuss these two different aspects in turn.

*Particle model of yeast growth:* Aside from being the principal model system used by cell biologists for elucidation of the genetic and biochemical factors responsible for regulating the eukaryotic cell cycle **[Mitchison 1971; Hartwell 1974; Forsburg and Nurse, 1991]** *S. cerevisiae* has also played a key role as the experimental focus of biophysical cell modelling studies due to its reproducible growth patterns and ease of assignment of distinct growth states under both light and scanning electron microscopes^22^ **[Hartwell and Unger, 1977; Chant and Pringle, 1995; Snijders and Pelkman, 2011; Cerulus et al. 2016; Mayhew et al. 2017]**. An important distinction to make at the outset of any discussion of cell modelling is that yeast can either be cultured in a liquid growth medium or on a solid growth medium (such as an agar plate) **[Andrews et al. 2016]**. In a well stirred liquid-growth medium the yeast tend to dissociate upon division, hence removing any associated positional considerations **[Hartwell and Unger, 1977; Lord and Wheals, 1980]** allowing them to grow unhindered until either resources become limiting or growth is slowed due to the release of quorum sensing factors at high yeast densities **[Andrews et al. 2016]**. Due to this simplifying feature, a lot of the early quantitative studies of yeast growth were caried out in liquid culture. Adopting the same descriptive transitive states of cell growth and division as shown in Fig. 2, Hartwell and Unger **(1977)** used experimental data gathered from analysis of such liquid culture systems to parameterize yeast growth rate constants under numerous growth conditions. Their quantitative modelling approach, restricted to the time domain and based on the assumption of exponential growth, yielded a number of important analytical forms relating differences in daughter and mother cell doubling times to overall growth rate **[Hartwell and Unger, 1977; Lord and Wheals, 1980]**. Recent quantitative studies of yeast growth have focused more carefully on these time constants by characterizing the dependencies of observable physical markers of cell growth on the different states of the division cycle **[Soifer et al. 2016; Mayhew et al. 2017]** whilst also examining the effect of noise and lineage on the stability of these time constants within a growing population **[Cerulus et al. 2016]**. We have tried to implement both the older classical viewpoints and these newer findings via combining routines involving the stochastic sampling of time constants (**Eqn. 1 and 2b**) with allowance for variability within and between yeasts according to their generation and lineage (**Eqn. 3 and 4**). For the sake of tractability, we sacrificed some observed mechanistic features of yeast growth such as the slow continual growth of mother cells to form slightly larger mother cells **[Vanoni et al. 1983]**. When yeast is grown on a solid culture (or even when grown in an unstirred solution) density and position effects will start to become a non-negligible aspect in the determination of yeast proliferation **[Shah et al. 2007; Rivas et al. 2014]**. The particle level description in MIL-CELL simulates yeast growth and division in two-dimensional culture as would be the case for yeast grown under restrictive conditions **[Zhao et al. 2018; Huberts et al. 2013]**. If growth is not restricted by use of a distance regulated coverslip arrangement, yeast will tend to form three dimensional colonies with observation of the underlying cells occluded by their placement within the colony **[Vulin et al. 2014; Ruusuvuori et al. 2014]**. Whilst the current approach could be quite simply extended to three-dimensions the primary^23^ reason for limiting it to two-dimensions is due to this inherent observational barrier associated with three-dimensional culture. Our approach for factoring in density effects is based on stochastic sampling against a pseudo-energy function (depicting the effort required to ‘push’ surrounding cells out of the way in order for internal cells to themselves grow or alternatively give birth to a new daughter cell) **[Eqn. 2c and d]**. As our yeast growth model is based on a set of rules it has characteristics of agent-based models first employed by Eden in the description of cell colony growth **[Eden 1960]**. Similar agent-based modeling approaches have been used to describe fungal growth **[Laszlo and Silman, 1993]**, confluent growth **[Lee et al. 1995]**, bacterial growth **[Kreft et al. 1998]**, tumor biology **[Drasdo and Höhme, 2005]** and nutrient limitation in two-dimensional yeast colony formation **[Banawarth-Kuhn et al. 2020]**. Whilst a defect of our model is the requirement for a fixed geometrical dependence (involving modelling mother cells as rectangular solids and daughter cells as spheres) to the best of our knowledge the approach specified in this paper is the only one capable of transitioning from locally confluent to non-confluent growth via specification of a single parameter (ε in Eqn. 2d – see Fig. 7). Without this ability growth will nearly always occur at the colony edge. Finally, by implementing a consistent color screen capable of accommodating nonlinear variation, the model has potential for provision of insight into questions relating to weak vs. strong phenotypes **(Eqn. 13, Fig. 5 – see Fig. 9 and 10 for examples) [Sharma and Liebman, 2012]**.

*Models of prion growth and transfer:* From the time of the initial association of prions with diseases such as Kuru and Scrapie **[Poser, 2002a, 2002b; Liberski, 2012]** there has been a great effort to quantitatively model both prion chemical and epidemiological dynamics **[Nowak et al. 1999]**. The first mathematical insight into polymer-based prion behavior was by Griffith **[1967]**. During a period of scientific uncertainty as to the exact biological nature of prions **[Gajdusek, 1977; Prusiner 1982; Weissmann 1991]**, the required chemical mechanisms and mathematical forms of various types of protein-based prion models were debated **[Come et al. 1993; Eigen, 1996; Nowak et al. 1999]**. Borrowing heavily from quantitative models applied to the polymerization of proteins such as hemoglobin, actin and tubulin polymerization **[Oosawa and Kasai, 1962; Wegner and Engel 1975, Hoffrichter et al., 1975; Oosawa and Asakura, 1975; Bishop and Ferrone, 1984; Flyvberg et al 1996; Hall, 2003; Hall and Minton, 2002, 2004]** early kinetic models of amyloid prion biology attempted to describe the spontaneous formation and differential transmission between host and recipient in terms of equivalent one-dimensional crystal growth and crystal seeding experiments [**Nowak et al. 1999; Masel et al. 1999; Pallito and Murphy, 2001; Craft et al. 2002; Hall and Edskes, 2004, 2009, 2012; Matthäus, 2006]**. With specific regard to the transmission of amyloid prions in yeast, three general types of approach have been attempted, (i) probabilistic models based on stochastic parameters **[Eaglestone et al. 2000; Cole et al. 2004; Byrne et al. 2009]**, (ii) kinetic models based on impulsive differential equations **[Lemarre et al. 2020]** and (iii) models based on spatial continuum dynamics of aggregate growth and movement **[Heydari et al. 2021]**.

In order to more clearly contrast the relative merits of these three alternative types of modelling approaches for describing prion growth in yeast against the approach adopted in the current work we first point out some of the distinctive points of the methods implemented within MIL-CELL for modelling amyloid growth and transfer. A strong point of the MIL-CELL method is the numerical approach employed for coupling the ordinary differential equation sets with the necessary partial differential equation forms required under conditions of changing volume and time **(Eqn. 11)**. The importance of including such concepts can be gathered from noting the predicted decrease in amyloid concentration during periods of rapid daughter cell growth with concomitant recovery of monomer concentration (due to it not being sequestered into amyloid - see **Fig. 9 and 10**). This numerical approach also has the added benefit of allowing for the direct usage of amyloid rate models determined and parameterized from quantitative experimental observations made under the typical constant volume in vitro conditions such as would be achieved using a microplate or cuvette system **[e.g. Xue et al. 2008; Hall et al. 2016]**. The kinetic models implemented in MIL-CELL are cast in terms of experimentally observed mechanisms previously demonstrated to have relevance to biology (e.g. variable internal versus endwise amyloid breakage relationships **[Hall, 2020]**, various nucleation and growth relationships **[Nowak et al. 1999; Pallitto and Murphy, 2001; Hall and Edskes, 2004, 2009, 2012; Hall and Hirota, 2009; Hirota et al. 2019]** and various higher order (end-to-end or lateral joining) amyloid association **[Zhao et al. 2016]**. Another important and distinctive feature of the current work is the specifiable component partition rate between mother and daughter cells during cell division **(Eqn. 10)**. Through inclusion of this term we have highlighted the need for its subsequent experimental or computational determination and/or further functionalization in relation to its size or yeast properties will likely prove key in understanding the generation time versus physical time disparities associated with analysis of yeast prion curing curves **[Marchante et al. 2017; Heydari et al. 2021]**.

In relation to the above description of MIL-CELL we note that the formulation of amyloid growth and transmission in the probabilistic models employed by the Cox, Morgan and Tuite collective **[Eaglestone et al. 2000; Cole et al. 2004; Byrne et al. 2009]** rely on a series of discontinuous decision-based stochastic jumps between yeast generations. Whilst this approach has an, in principle, capability of monitoring the spread of the yeast prions with yeast position information in practice this was not employed by the authors^24^ **[Byrne et al. 2009]**. An advantage of the probabilistic approach is its ready usage in fitting data gathered from experimental curing curves **[Byrne et al. 2009]** (**e.g. see Fig. 10**) however without any continuous physical governance of chemical behavior this type of model is very much limited by the veracity of the assumptions governing the component behavior and transfer between generation time points. A different approach for describing amyloid growth and transmission in yeast, was based on the use of sets of impulsive ordinary differential equations [**LeMarre et al. 2020].** Although the modelling approach was decoupled from the physical placement of the yeast on the plate the authors used this method to demonstrate existence of a bistable regime corresponding to the possible coexistence of [PSI+] and [psi−] within the same colony – the so-called colony sectoring experiment **[Sharma and Liebman, 2012; LeMarre et al. 2020]**. Some negative aspects of the method adopted by LeMarre et al. include its slightly unphysical aggregation mechanism, its reliance on a fixed cell division time and the use of set rules for partition of cellular contents made on the basis of mother and daughter volumes alone. One further weak point is that the formulation of the differential equation set seems unsuited to conditions involving both changes in volume and time. The final alternative approach which we discuss here is the use of spatial continuum dynamics for describing aggregate growth and transfer **[Heydari et al. 2021]**. Based on realistic descriptions of yeast geometry and internal components this approach is potentially superior (although much more computationally intensive) to the one described in the present work, however at present it has only been applied to description of a protein monomer-dimer interaction within a single dividing cell **[Heydari et al. 2021]**. Also noted by the authors, the continuum dynamics approach potentially breaks down at low absolute molecular number, potentially necessitating a switch to a discrete particle simulation method such as the Brownian dynamics approach **[Hall et al. 2006; Auer et al. 2006; Hall and Hoshino, 2012]**.

### (ii)#Biochemical complexity of the epigenetic phenomenon

The starting motivation for the MIL CELL project was to provide a means for modelling the nonlinear dynamics of prion-based epigenetic inheritance in yeast. The term ‘molecular epigenetics’ is frequently understood as referring to the differential transfer of an active biochemical factor between mother and daughter cells such that the biochemical factor is capable of influencing recorded expression profiles in a manner not necessarily consistent with the genetic sequence information contained within the chromosomal DNA [**Waddington 1942; Manjrekar 2017**]. Two early realizations of such an ‘active biochemical factor’ for affecting changes in gene expression included (i) chromosome specific DNA methylation **[Rhazin and Riggs, 1980; Weissbach, 2013]**, and (ii) post-translational modification of the histone proteins in chromatin **[Burggren 2016; Manjrekar 2017; O’Kane and Hyland, 2019]**. In normal mitotic cell division, the distribution of genetical material is effectively digital in nature, with one copy retained by the mother and one copy transferred to the daughter. However, if an epigenetic factor is not evenly distributed between the, in principle, identical segregated genetic material, there will in effect be an unequal transmission of genetic material between mother and daughter cells which can lead to differences between them **[Weissbach, 2013; O’Kane and Hyland, 2019]**. Similarly, for the case of meiotic cellular division with subsequent sexual reproduction, such unequal distribution of epigenetic factors amongst the gametes can significantly affect the likelihood of observing normal ‘expected’ Mendelian phenotypes on the basis of genotype [**Kota and Feil, 2010**]. Another important, yet different, area of epigenetics arises from the maternal effect – a catch all designation used to describe the unequal sharing (asymmetric division) of soluble cytosolic components between the mother and daughter cells [**St Johnston 1995; Bonasio et al. 2010**]. The unequal aspect of sharing may be due to simple stochasticity (when the absolute number of the components is sufficiently low to allow differences to arise from statistical chance) **[Bonasio et al. 2010; Cerulus et al. 2016]** or from specialist biological mechanisms that either preferentially retain damaged components [**St Johnston, 1995; McFaline-Figueroa et al. 2011; Yange et al. 2015**] or preferentially promote the uptake of advantageous or required components able to facilitate the best possible outcome for the nascent daughter cell [**McFaline-Figueroa et al. 2011**]. The chemical lifetime of these added components has dramatic consequences for their ability to act as trans-generational epigenetic factors **[Fitz-James and Cavalli, 2022]**. As shown from microinjection experiments, whilst relatively short-lived components, such as various coding/noncoding RNA or enzymes can influence the immediate growth behaviour of the daughter cell^25^, they do not necessarily show particularly strong genetic linkages to subsequent generations **[Lim and Brunet, 2013; Fitz-James and Cavalli, 2022]**. However, long-lived, structurally persistent states, able to perpetuate and replicate themselves over the time-course of the cell-division cycle, are themselves able to be inherited and are therefore capable of showing strong epigenetic linkage patterns^26^ **[Wickner et al. 2015]**. In both yeast, and bacteria, the original prototypic cytosol based epigenetic factors were small pieces of circularized nucleic acid known as plasmids [**Wickner and Leibowitz, 1977; Gunge, 1983**]. Acquisition or loss of a plasmid was shown to confer additional traits such as antibiotic resistance or sexual mating preference [**Gunge, 1983**].

Somewhat more recently, a second class of cytosolic epigenetic factor comprised of amyloid polymer has been found to be common in certain yeast and mold species **[Wickner 1994; Wickner et al. 1995; Wickner et al. 1997; Dos Reis et al. 2002; Tuite and Serio, 2010; Halfman et al. 2012; Wickner et al. 2020]**. Referred to as yeast (and fungal) prions ^27^ , these epigenetic components are physically constituted by structurally persistent amyloid homopolymers and due to their ability to effect a phenotype, are sometimes referred to as protein genes [**Wickner et al. 2015**]. The question as to whether or not these new classes of amyloid-based epigenetic components act to improve organism fitness by playing a positive role **[Halfman et al. 2012; Garcia and Jarosz, 2014; Wang et al. 2017]**, decrease organism fitness thereby acting as a disease **[Wickner et al. 2011]**, or even constitute an as yet unknown biochemical function, is still an open one **[Tuite and Serio, 2010]**. In our model we left this question open by parameterizing the free concentration of amino acids in terms of the total build-up of monomer within prion form **(Eqn. 6)**. Growth rates could be similarly parametrized in terms of either total prion levels or the amount of prion of a particular size **[Hall and Edskes, 2009]**. However, despite an ongoing debate over the role played by prions, the widespread existence of a range of different types of yeast prions, and the biomolecular components that interact with them, has been concretely established **[Wickner et al. 2015; Wickner et al. 2020]**. Aside from the [PSI+], [URE2] and [RNQ1] prion elements already discussed in this work there has been approximately ten other types of yeast prions discovered (with each prion based on a different amyloid protein component) **[Wickner et al. 2015]**. Alongside research on the prion components themselves, has been the discovery of biochemical components that interact with prions to modulate their behaviour to achieve the following functional outcomes (i) prion re- solubilization, (ii) prion degradation, (iii) prion selective segregation and (iv) prion sequestration **[Wickner et al. 2015]**. Within yeast, a range of protein regulatory subsystems have been shown to be active in these different forms of prion modulation with a non-exhaustive list including the following; chaperone systems **[Verghese et al. 2012; Chernova et al. 2017]**, ubiquitin-proteasome system **[Berner et al. 2018]**, autophagy system **[Suzuki and Ohsumi, 2007]**, aggresome systems **[Miller et al. 2015]**; vacuoles (the yeast lysosome) **[Armstrong, 2010]**; system for asymmetric segregation of damaged proteins **[Coelho and Tolić, 2015]**, GET pathway proteins [**Borgese and Fasana, 2011**] and the Btn2-Cur1 system **[Wickner et al. 2014]**. Interestingly, MIL-CELL offers the potential to replicate the functional outcome from the up or down regulation of these various prion-regulating components through specification and parameterization of the governing model constants.

### (iii)#Potential wider application of MIL-CELL to disease and non-disease cellular processes

In creating MIL-CELL the motivation was to develop an easy-to-use tool capable of shedding light on the phenomenon of epigenetic inheritance in the budding yeast Saccharomyces cerevisiae associated with amyloid- prions **[Wickner, 1994; Wickner et al. 1995, 1997; Tuite and Serio, 2010; Wickner et al. 2020]**. However, due to the generality of the modelling approach used to describe both cell growth/division, and the chemical behavior of the cytosolic component, MIL-CELL has potentially significant capabilities to provide insight into a range of other cellular phenomenon. In this section we discuss some of these additional capabilities in terms of MIL-CELL’s potential use for casting light on disease and non-disease cellular processes.

*Using MIL-CELL to investigate disease at the cellular level:* Although not limited to the following we discuss MIL- CELL’s applicability to, amyloidosis, cancer and mitochondrial dysfunction. **(i) Amyloidosis:** As of 2022 there are 42 proteins known to form amyloid in humans **[Buxbaum et al. 2022]**. The various amyloidosis diseases (including Alzheimer’s disease, Type 2 diabetes and cardiac amyloidosis) all involve the formation of significant amounts of amyloid aggregates which interact negatively with tissue due to either aggregates possessing an intrinsic cytotoxicity, or through physical effects manifested from amyloid infiltration into the tissue space, changing its material properties and diminishing its normal function **[Hardy and Higgins, 1992; Merlini and Belloti, 2003; Hall and Edskes, 2009, 2012; Martinex-Naharro et al. 2018; Fornari et al. 2019]**. MIL-CELL could effectively replicate such empirical experimental realities by including export of monomer from a cell to the interstitial space with subsequent description of its diffusion and aggregation within that space given by use of either compartment modelling **[Craft et al. 2002]** or sets of partial differential equations reflecting two-dimensional diffusion reaction equations **[Matthäus, 2006]**. Cell growth rates and death could be made functions of the extent and length of exposure to amyloid in the interstitial space. Although the amyloidosis diseases were originally characterized by the extracellular formation and deposition of amyloid [**Buxbaum et al. 2022**] the last thirty years has seen both a growing recognition of the intracellular accumulation and processing of amyloid in diseases such as Alzheimer’s **[Hardy and Higgins, 1992; Glabe, 2001]** via both intracellular aggregation mechanisms and endosomal and lysosomal transport of external amyloid **[Bayer and Wirths, 2010]**. Also, there are a number of amyloidosis-related disease states which primarily involve intracellular accumulation of amyloid aggregates, such as Alzheimer’s related tau amyloidosis **[Nizynski et al. 2017]**, Parkinson’s disease related α-synuclein amyloid formation **[Lücking, 2000]** and Huntington’s disease related huntingtin amyloid formation **[Ross and Tabrizi, 2011]**. In its present state MIL- CELL could be used to investigate the effect of various kinetic mechanisms on the time course and development of amyloid within a cell space (e.g. see **Fig. 4**) and tying the build-up of amyloid beyond a certain level to cell toxicity and cell death. With very slight extension MIL-CELL could also be adapted to feature amyloid transfer between cells via mechanisms dependent on either cell rupture following death or transfer via exosome formation **[Steiner et al. 2011]**. **(ii) Cancer:** Due to its use of a particle model for cell growth and division MIL-CELL holds significant potential for use in the investigation of the two basic sub-fields of cancer biology described as initiation and migration **[Bertram 2000; Riggi et al. 2018]**. Initiation of cancer involves the flipping of one of a large number of genetic/biochemical switches which transforms a previously healthy cell (having defined growth and division patterns and specific cellular morphology) to a cancerous cell which typically lacks tight control of growth and division and loses its normal cell morphology and respect of local confluence **[Bertram 2000]**. Such a situation could be reconciled within MIL-CELL by individually assigning cells a local confluence parameter ε **(Eqn. 2c, d)** (with the value determined by either a stochastic incidence, a defined neighbor relationship or position within a colony or the build up of a chemical component) and local growth rates and local variability terms **(Eqn. 1, 2a, b, 3, 4)**. A hallmark of transformed cancer cells is their tendency to undergo migration and spread through the body in a process known as metastasis **[Riggi et al. 2018]**. Their departure from tissue of origin to the bloodstream (intravasation) and their movement from the bloodstream into a new tissue (extravasation) involves purposeful migration through tissue with the necessary displacement of surrounding cells **[Riggi et al. 2018]**. Directed motion within MIL-CELL could be simply implemented by assigning a semi-random drift velocity to a transformed cancer cell growing within the colony. **(iii) Mitochondrial dysfunction:** Existing within every eukaryotic cell, mitochondria are semi-autonomous organelles that contain their own genome and which coordinate their growth and replication with that of their host cell **[Ernster and Schatz, 1981; Bock and tait, 2020]**. Mitochondria are largely responsible for the production of ATP (Adenosine Tri-Phosphate) within every eukaryotic cell by virtue of the fact that they contain the biochemical machinery necessary for carrying out the three essential metabolic processes known as the Tricarboxylic Acid Cycle (TCA – breakdown of citric acid) **[Ernster and Schatz, 1981]**, β-oxidation (breakdown of fatty acids) **[Ernster and Schatz, 1981]** and oxidative phosphorylation (coupling the oxidation of high energy reduced cofactor (NADH - nicotinamide adenine dinucleotide) via oxygen to form water and oxidized cofactor (NAD+) with the production of 3 ATP) **[Ernster and Schatz, 1981].** Yeast cells possess a small number of mitochondria (∼10) **[Vevea et al. 2014]**, mid-sized mammalian eukaryotic cells can contain hundreds to thousands **[Dewey and Fuhr, 1976]** whilst large cells, such as neurons, can contain thousands to millions **[Misgeld and Schwarz, 2017]**. Some forms of intracellular amyloid have been observed to both directly, and indirectly, damage mitochondria [**Bayer and Wirths, 2010].** Additionally, asymmetric transfer of mitochondria during cell division can result in significant differences in growth rate between mother and daughter cells as well as contributing to numerous diseases (such as cancer and also, in a somewhat circular fashion, amyloidosis, amongst many others) **[Annelsey and Fisher, 2019]**. In its current state MIL-CELL has the ability to mimic mitochondrial replication inside the cell and transfer within a dividing cell population (by specifying the initial number of mitochondria and setting f_G_ > 0, (k_A1_)_αβ_ > 0 and f_N_ = f_A1_ = f_A2_ = 0) and also by tying cell growth rate to mitochondrial number). Further developments could involve explicit specification of mitochondrial damage via amyloid accumulation and preferential retainment/donation of such damaged mitochondria during cell division.

*Using MIL-CELL to investigate non-disease processes at the cellular level:* As a model of cell growth MIL-CELL holds potential for investigating a range of fundamental aspects of cell biology not necessarily associated with disease. Without restriction we introduce three open questions in cell biology to which MIL-CELL could provide useful insight, namely cell variability within a clonal population, temporal and morphological heterogeneity in cellular differentiation pathways and differences in mechanisms of cell death dependent on generational versus linear aging. **(i) Cell variability:** Cultured cells are often used as the first type of *in vivo* model for testing efficacy of a potential drug, the cytotoxicity or mutagenicity of a dangerous substance, or as a production platform in the creation of a useful biochemical **[Stevens and Baker, 2009; Hong et al. 2018]**. To eliminate sources of variability, the test culture is typically produced via dilutional plating to ensure a single clonal population **[Hong et al. 2018]**. Despite the existence of an isogenic population the individual cells within the culture will exhibit variation not just in fundamental observable traits, such as mRNA and protein expression, cell size and morphology and cell growth and division rates (to name but a few) **[Stockholm et al. 2007]**, but will also exhibit variation in response to the drug or dangerous compound being tested **[Moore et al. 2018]**. Knowledge of the functional form of this variation and how it evolves over time is necessary for assessing confidence in experimental results **[Stockholm et al. 2007; Moore et al. 2018; Hong et al. 2018]**. MIL-CELL features a novel implementation of variability of cell growth rate constants based on random sampling from a distribution produced by recursive formula updated with each cell generation (**Eqn. 3 and 4**). It will prove interesting to see how accurately this formulation can replicate variability recorded from microscopy or single cell cytometry studies [**Stockholm et al. 2007; Moore et al. 2018**]. **(ii) Temporal and morphological heterogeneity in cellular differentiation pathways:** Developmental biology, such as is typified by the production of an entire organism from a single fertilized egg, is a process that requires exquisite spatial and temporal control of cell the division and differentiation pathways as well as extremely robust mechanisms for dealing with environmental variation at different stages of the developmental process **[Oates et al. 2009]**. Although originally cast in terms of a culture of single cell organisms one obvious extension of the MIL-CELL program would be to allow for both cell differentiation and cell-to-cell association to provide a primitive model of tissue formation **[Keller, 2013]**. Such programmed differentiation could be age based (either chronological or replicational) or position based (e.g. determined in relation to time spent at the interior or edge of the culture **/** proximity to a new class of differentiated cell) or the cell’s location within a gradient of externally derived environmental signals **[Oates et al. 2009; Keller, 2013]**). **A priori** developmental programming combined with rapid simulation within MIL-CELL offers the prospect of identifying stable robust developmental strategies. **(iii) Cell death:** In a multicellular organism, regulation of the process of cell death is an absolute requirement for both its correct development and continued maintenance/perpetuation **[Doherty and Baehrecke, 2018]**. Cell death can occur in one of three general ways, apoptosis (purposeful breakdown of the genetic material), autophagy (literally ‘self-eating’ due to the formation of a large internal double membraned vacuolar body known as the autophagosome which engulfs and transports cellular contents to the lysosome for subsequent breakdown) and necrosis (cell death resulting from irreparable cellular damage caused by injury or disease) **[Doherty and Baehrecke, 2018]** with the first two of these considered as alternative methods of programmed cell death **[Fuchs and Steller, 2011; Doherty and Baehrecke, 2018]**. The relationship between these three types of cell death to both the age of a cell (chronological and replicational) and its exposure to an external or internal signal is an area rich in potential for investigation and MIL-CELL has much of the required mathematical formulation in place to provide insight into these types of study topics.

## Conclusions

The increasing availability of computing power holds significant potential to aid with the interpretation of difficult to understand experiments of complex biological phenomena [e.g. **Cerulus et al. 2016; Hall, 2020; LeMarre et al. 2020**]. Effectively established by Wickner in the early 1990s, the sub-field of epigenetic inheritance conferred by yeast amyloid prion growth and transmission within a dividing population of yeast cells is a particularly mature example of such complex biological phenomena [**Wickner 1995**]. By coupling a multiscale dynamic model of cell growth and amyloid kinetics together with the powers of a relational database, the MIL-CELL program described within the current paper, will prove useful in the interpretation of the results of experiments involving amyloid transmission between dividing cells. Recently, the asymmetric transmission of both amyloid, and other cytosolic components, has been understood as holding importance across many areas of cell biology, from fields such as epigenetics (as discussed in this work) to cellular differentiation, cellular aging and death and the study of diseases such as cancer, mitochondrial dysfunction and amyloidosis. We hope that the MIL-CELL program may also assist in shedding light on these additional topics in the future.

## Acknowledgements

DH would like to sincerely thank Profs. A.S. Foster, J. Harris, H. Edskes, M. Okada, T. Sumikama, H. Flechsig, C. Franz, K. Nagayama, Reed Wickner and H. Nakamura for a series of helpful written comments on an earlier draft of this work. DH would also like to thank Dr. Reed Wickner and Dr. Daniel Masison (and their associated laboratory members) for helpful discussion at the outset of this work. DH acknowledges funding associated with the receipt of a ‘Tokunin’ Assistant Professorship carried out at the WPI-Center for Nano Life Science, Kanazawa University. This work was supported, in part, by KAKENHI Start-Up grant 21K20633, and WPI ‘Funds in Aid of Research’ grant awarded to D.H.

## Conflict of Interest Statement

D.H. reports no conflict of interest.

^1^ And indeed from higher ploidy states.

^2^ Acting in this role, budding yeast helped to extend our quantitative understanding of molecular biology initiated by Delbruck and colleagues’ studies on viruses and bacteria [**Kay, 1985; Morange 2000**].

^3^ The term epigenetics was first coined by Waddington in 1942 as he attempted to jointly study the relatively new fields of embryology and genetics (epigenetics = ‘**epi**genesis’ + ‘**genetics**’) [**Waddington, 1942**]. A simple operational definition of an epigenetic factor is something that can effect a phenotypic trait (that can be passed on to offspring) but which is not simply specified by a linear chromosomal gene sequence of the parent. This operational description is inclusive of the two standard definitions of epigenetic factors discussed more carefully by Haig [**Haig, 2004**]. Molecular epigenetic factors can be considered as trans or cis. A ‘trans’ epigenetic molecular factor is transmitted via partitioning of the cytosol during cell division, whereas ‘cis’ epigenetic molecular factors are physically associated with the chromosomal DNA and are passed on via chromosomal segregation during cell division [**Bonasio et al. 2010**].

^4^ Although mentioned here due to the fact that it is a common epigenetic factor, DNA methylation does not occur to an appreciable extent in yeast [**Tang et al. 2012**].

^5^ The normal function of Ure2p is that of a negative regulator of nitrogen catabolism [**Courchesne and Magasanik, 1988; Coschigano and Magasanik, 1991**].

^6^ Carried out by growth of yeast in media containing/not containing limited amounts of an alternative nitrogen source [**Lacroute 1977; Wickner et al. 1995**].

^7^ The normal function of Sup35 is as a release factor in the multi-component translation termination complex [**Didichenko et al. 1991; Tuite, 1995**].

^8^ Yeast can be engineered to contain genes with a premature translation stop codon. Due to Sup35 role as a translation arrest factor [**Didichenko et al. 1991; Stansfield et al. 1995**] the proteins coded by such genes with a premature stop codon can only be expressed when Sup35 is non-functional, such as is the case when Sup35 exists as an insoluble inactive component within an amyloid prion. Using this reporter mechanism a range of functional markers are possible with the standard involving an ADE1 gene mutant containing a premature stop codon. The functional enzyme produced from the ADE1 gene catalyses the enzymatic cleavage of a red coloured intermediate in the adenine biosynthetic pathway. Presence of the functional Sup35 monomer prevents expression of the ADE1 gene product thereby affording the yeast a red color and necessitating that the yeast be grown in media containing the adenine (see **Fig. 5**).

^9^ The yeast vacuole system is the equivalent of the lysosome system in mammalian eukaryotic cells.

^10^ At a time sufficiently progressed that the likelihood of occurrence of the kinetic transition at zero local density is one i.e. P(I→J,ρ_local_ =0,t®¥) = 1.

^11^ We have adopted this latter physical viewpoint of the resistance to insertion/growth to yeast segments i.e. the primary resistance is generated by pushing other yeast out of the way due to either their interaction with the surface or fluid surrounds. ^12^ By reduced we mean 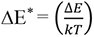 with k being the Boltzmann constant and T the absolute temperature. We acknowledge that the Boltzmann distribution may not be the most appropriate term due to the macroscopic nature of the particles involved (e.g. cells). However, even if applied imperfectly, the selection process is well formulated and its consistent application is sufficient to provide insight into the internal vs external growth of cells within or around the yeast colony.

^13^ Yeast movement is broken down into the positive ┴ and negative ┬ directions perpendicular to the plane of insertion with the total number of yeast segments having to be ‘pushed’ in these respective directions denoted as N_(┴)_ and N_(┬)_. If N_(┴)_ = N_(┬)_ a 50:50 random split is made between moving the N_(┴)_ or N_(┬)_ set. In the case that the yeast growth is effectively fixed in place on the plate i.e. ε → ∞ then the Metropolis selection criteria reduces to one of yeast particle non-overlap, a situation requiring that the distance, d_mn_, between any two yeast segment centers*, C_m_(x,y,z) and C_n_(x,y,z), is larger than the sum of their respective radii, R_m_ and R_n_, 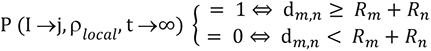. (*To make the computation easier each mature yeast is considered as containing two spherical segments within the rectangular solid of aspect ratio 2).

^14^ An alternative point of view is that it is the loss of functional monomer that causes a deleterious loss of function.

^15^ We will speculate on the mechanism of the self-limitation in the discussion.

^16^ For simplicity we set n = 2. This simplifying assumption has been discussed elsewhere **[Hirota et al. 2019]**.

^17^ This is known as the uni-directional monomer addition, bi-directional monomer loss assumption for which there is significant experimental support [**Heldt et al. 2011; Beun et al. 2016**].

^18^ The nascent daughter is referred to as the ‘bud’ prior to septum closure.

^20^ Bearing on mind that we are solely considering monolayer growth in this paper in the form of yeast growing in a two- dimensional restricted space.

^21^ In practice yeast are plated on media containing guanidine HCl which prevents Hsp104-based cleavage of amyloid prions. Single colonies are selected and then replated media containing media lacking adenine, ensuring that only yeast containing amyloid prions will be able to grow.

^22^ SEM measurements can reveal the bud scars in mother yeast cells produced at each cell division **[Chant and Pringle, 1995]**. Bud scars can also be visualized by fluorescence microscopy upon staining yeast with calcofluor white.

^23^ Another problem is the differential transport and competition for solutes from the growth medium that is associated with three-dimensional cell growth **[Vulin et al. 2014]**.

^24^ Despite developing a perhaps superior digital indexing system to the one described in the current paper.

^25^ In the role of an initial condition.

^26^ When they play some positive or negative role in cell growth and can therefore produce an observable phenotype.

^27^ The prion term was first coined by Stanley Prusiner who discovered the amyloid-based molecular origins of set of closely related disease spongiform encephalopathies that are characterized by chronic brain wasting, dementia and eventually death (Scrapie in sheep, BSE (bovine spongiform encephalopathy) in cows, and Kuru and variant CJD (Creutzfeld-Jakob disease) in humans (amongst others) [**Prusiner 1982**].

