## Supplementary for "MIL Cell – A tool for multi-scale simulation of yeast replication and prion transmission"

**Supplementary File 1 - Damien Hall**

The MIL-CELL program can be downloaded freely using the following ftp link.

https://drive.google.com/drive/folders/1xNBSL_2sGNkyXfYLYUyXjyM9ibGAcQUL?usp=sharing

By downloading the material you are agreeing to the following academic end user license agreement (EULA)

**MIL-CELL User Agreement**

This document is an End User License Agreement (EULA) for the MIL-CELL software operating between the downloading party and the author of this software (Damien Hall).

(1) The academic version of this software is written in MATLAB CODE R2020 programming language and packaged as an executable file compatible with a windows 10 operating system.

(2) While every care has been taken to make sure that the MIL-CELL software works correctly the user is provided with no guarantee of the correctness of the results produced. These programs are provided as is and are to be operated at the risk of the user. The user also assumes all risk associated with any damage that these programs may cause to the computers being used to run these programs and the data stored within these computers.

(3) These programs are provided freely for academic use only to the downloading user. The user may not subsequently claim authorship of these modified programs or subsequently host, post or independently distribute these programs to a third party.

(4) These programs will be updated with some additional functionality over time. If you would like to be alerted of these additions/improvements via email under the identical terms of this current agreement please send your name and email to with 'MILL-CELL USER GROUP REQUEST' indicated in the subject line.

(5) When using these programs for publication please cite the reference Hall, D. (2022). MIL Cell – A tool for multi-scale simulation of yeast replication and prion transmission. BIORXIV 2023

(6) By providing this program to the downloading party for academic use, the author does not relinquish any present or future intellectual property rights to the MIL-CELL software suite.
